# Drug Reinforcement Impairs Cognitive Flexibility by Inhibiting Striatal Cholinergic Neurons

**DOI:** 10.1101/2022.10.27.514125

**Authors:** Himanshu Gangal, Xueyi Xie, Yifeng Cheng, Xuehua Wang, Jiayi Lu, Xiaowen Zhuang, Amanda Essoh, Yufei Huang, Laura N. Smith, Rachel J. Smith, Jun Wang

## Abstract

The mechanisms underlying the reduction in cognitive flexibility associated with reinforcement of addictive substance use are unknown. This reinforcement is mediated by substance-induced synaptic plasticity in direct-pathway medium spiny neurons (dMSNs) that project to the substantia nigra (SNr). Cognitive flexibility is mediated by cholinergic interneurons (CINs), which receive extensive local inhibition from the striatum. Here, we report that cocaine or alcohol administration caused a long-lasting potentiation of local inhibitory dMSN→CIN transmission in the dorsomedial striatum (DMS), a brain region critical for goal-directed behavior and cognitive flexibility. This dMSN→CIN potentiation reduced CIN firing activity. Furthermore, chemogenetic and time-locked optogenetic inhibition of DMS CINs suppressed cognitive flexibility in an instrumental reversal learning task. Importantly, rabies-mediated tracing and physiological studies revealed that SNr-projecting dMSNs, which mediate reinforcement, sent axonal collaterals to inhibit DMS CINs, which mediate flexibility. Our findings demonstrate that the local inhibitory dMSN→CIN circuit mediates a reinforcement-induced reduction in cognitive flexibility.

**HIGHLIGHTS:**

1. Cocaine reinforcement inhibits striatal cholinergic interneurons (CINs) and impairs cognitive flexibility.
2. Optogenetic and chemogenetic CIN inhibition impairs cognitive flexibility.
3. Reinforcement behaviors potentiate inhibitory transmission from direct-pathway medium spiny neurons (dMSNs) to CINs.
4. Substantia nigra-projecting dMSNs mediate reinforcement and also send collaterals that inhibit CINs.

## INTRODUCTION

Natural rewards and addictive substances such as cocaine and alcohol increase dopaminergic activity in the striatum, which mediates over 90% of voluntary behaviors. The elevated dopamine level facilitates corticostriatal synaptic plasticity in direct-pathway medium spiny neurons (dMSNs) (Everitt and Robbins, 2005; Luscher, 2016; Pascoli et al., 2018; Pascoli et al., 2015; Schultz, 1998; Shan et al., 2014). This plasticity then drives an animal to repeat actions that previously resulted in rewards, thus promoting reinforcement learning (Luscher and Janak, 2021; Luscher and Malenka, 2011; Luscher et al., 2020; Schultz, 1998). The projection of dMSNs to the substantia nigra (SNr) forms the direct pathway of the basal ganglia (Cheng et al., 2017; Gerfen and Surmeier, 2011; Kreitzer and Malenka, 2008), which is known to be particularly important for mediating reward-induced reinforcement (Hellard et al., 2019). Drug-free optogenetic studies have demonstrated that intracranial self-manipulation of dMSN→SNr activity alone is sufficient to induce reinforcement (Kravitz et al., 2012; Lalive et al., 2018), confirming the role of the direct pathway in this behavior. These findings suggest that positive reinforcers (e.g., natural rewards and addictive substances) generally potentiate dMSN activity, which then drives behavioral reinforcement.

Reinforcement of drug and alcohol use has also been associated with reduced cognitive flexibility, which contributes to compulsive drug and alcohol use (Barker et al., 2017; Izquierdo and Jentsch, 2012; Jentsch et al., 2002; Luscher *et al*., 2020; Ma et al., 2021; McCracken and Grace, 2013; West et al., 2021). However, the neurocircuitry involved in this reinforcement-induced inflexibility has not been identified. Cognitive flexibility facilitates adaptation to environmental changes in order to obtain a reward or avoid punishment (Ragozzino, 2007). In animal studies, cognitive flexibility can be measured using reversal learning tasks where animals are trained on a set of action-outcome contingencies that are then reversed; animals are considered flexible if they successfully learn the new association (Ragozzino, 2007). Substantial evidence indicates that a rise in striatal acetylcholine (ACh) is required for reversal learning (Aoki et al., 2015; Bradfield et al., 2013; Galaj et al., 2019; Matamales et al., 2016; Prado et al., 2017; Ragozzino, 2007; Ragozzino and Choi, 2004). Striatal ACh is mainly released by cholinergic interneurons (CINs). CINs primarily receive GABAergic inputs (Gonzales et al., 2013) and extensive local afferents from the striatum itself (Guo et al., 2015), which is composed of GABAergic dMSNs and indirect-pathway MSNs (iMSNs) (Cheng *et al*., 2017; Gerfen and Surmeier, 2011; Kreitzer and Malenka, 2008). Therefore, CINs are likely to receive inhibitory inputs from MSNs in the striatum. Given that drug reinforcement enhances striatal dMSN activity, we hypothesized that inhibitory dMSN→CIN transmission played an essential role in mediating drug-induced suppression of cognitive flexibility.

We observed that cocaine or alcohol exposure increased GABAergic inputs from dMSNs to CINs and suppressed CIN firing in the DMS, a brain region that regulates goal-directed behaviors and cognitive flexibility (Bradfield *et al*., 2013; Matamales *et al*., 2016). Chemo- or optogenetic inhibition of CINs reduced cognitive flexibility. Importantly, SNr-projecting dMSNs that mediate reinforcement also sent collaterals that inhibited CINs. Our data demonstrate that the inhibitory dMSN→CIN connection mediates the reinforcement-induced reduction in cognitive flexibility.

## RESULTS

### Cocaine reduces flexibility and inhibits CINs

To investigate how cocaine administration affects cognitive flexibility, we intraperitoneally (i.p.) injected C57BL/6 mice with cocaine or saline for 7 d (Pascoli et al., 2014; Pascoli et al., 2012) (Fig. 1a). Mice that received cocaine showed a significantly greater total distance traveled (Fig. 1b; *F*_(1,17)_ = 41.87, *p* < 0.001). Animals were then trained on a two action (A)-outcome (O) reversal learning task, where nose-poking on the left port (A1) resulted in the delivery of a food pellet (O1) to the left magazine (A1→O1), and nose-poking on the right port (A2) led to sucrose delivery (O2) in the right magazine (A2→O2) (Fig. 1c) (Bradfield *et al*., 2013; Matamales *et al*., 2016). This task has previously been used to study cognitive flexibility and reversal learning (Bradfield and Balleine, 2017; Bradfield *et al*., 2013; Matamales et al., 2020; Matamales *et al*., 2016). We trained the animals on these action-outcome contingencies for 11 consecutive sessions (Fig. 1c). Cocaine- and saline-administered mice exhibited no significant difference in total nose-pokes (Fig. 1d; *F*_(1,12)_ = 1.15, *p* > 0.05). Following these initial sessions, we reversed the action-outcome contingencies of the task (Fig. 1e): A1→O2 and A2→O1 (Bradfield *et al*., 2013; Matamales *et al*., 2016). We found significantly fewer nose-pokes by the cocaine-administered mice during these reversal sessions, as compared with the saline controls (Fig. 1f; *F*_(1,12)_ = 7.26, *p* < 0.05). These results indicate that although cocaine administration does not alter the initial learning of action-outcome contingencies, it impairs reversal learning and reduces cognitive flexibility.

**Figure 1.**
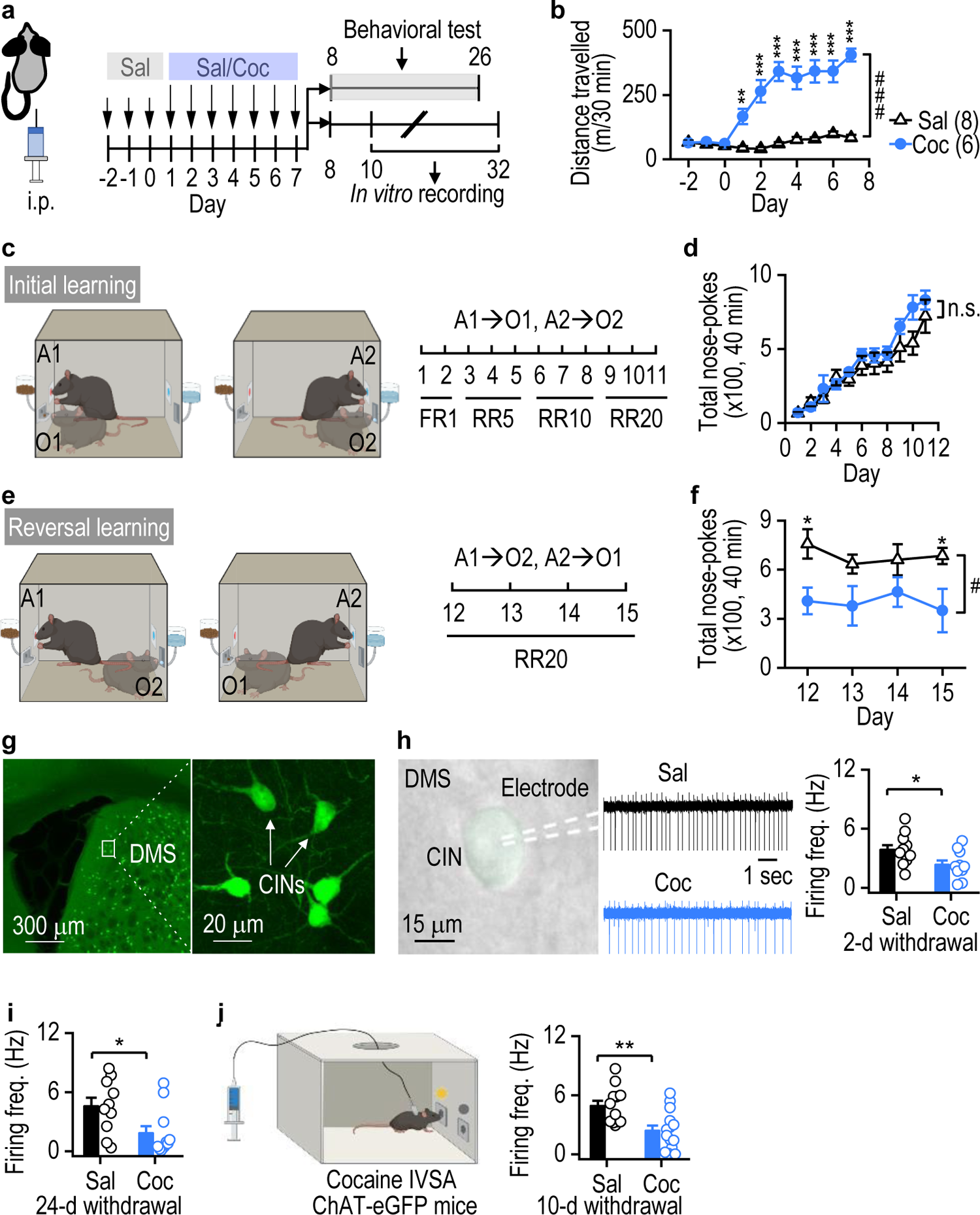
Cocaine administration compromises reversal learning and reduces DMS CIN activity. **a**, Schematic of intraperitoneal (i.p.) injections of saline (Sal) or cocaine (Coc, 15 mg/kg) and the subsequent behavioral or electrophysiological measurements. **b**, Repeated cocaine injections caused locomotor sensitization in C57BL/6 mice; ^###^*p* < 0.001 for the indicated comparison; \*\*\**p* < 0.001, \*\**p* < 0.01 versus the saline group on the same day. **c**, Initial instrumental action (A)-outcome (O) contingency training: mice nose-poked the left port (A1) to receive food pellets (O1; A1→O1) and the right port (A2) to receive sucrose solution (O2; A2→O2). FR1: fixed ratio 1; RR5-20: random ratio 5-20. **d**, Cocaine did not alter total nose-pokes during initial learning; n.s., not significant. **e**, Reversal training (A1→O2, A2→O1). **f**, Cocaine inhibited total nose-pokes during reversal learning; ^#^*p* < 0.05; \**p* < 0.05. **g**, Images of DMS CINs in a ChAT-eGFP mouse. **h**, Repeated cocaine injections reduced the spontaneous firing frequency of DMS CINs 2 d after the last injection; \**p* < 0.05, n = 11 neurons from 3 mice (11/3) (Sal) and 10/3 (Coc). **i**, Cocaine-induced inhibition of DMS CIN activity persisted 24 d after the last injection; \**p* < 0.05, n = 10/3 (Sal) and 11/4 (Coc). **j**, Cocaine intravenous self-administration (IVSA) also inhibited DMS CIN firing 10 d after the last exposure; \*\**p* < 0.01, n = 12/4 (Sal) and 14/4 (Coc). Two-way RM ANOVA followed by Tukey *post-hoc* test (b, d, f); unpaired *t* test (h-j).

Cognitive flexibility is known to be modulated by DMS CINs (Bradfield *et al*., 2013; Matamales *et al*., 2016). Therefore, we next investigated whether the same cocaine treatment altered DMS CIN activity in choline acetyltransferase (ChAT)-eGFP mice, in which CINs expressed GFP (Fig. 1a, 1g, 1h, Supplementary Fig. 1a). DMS slices were prepared 2 d after the last cocaine injection, and cell-attached recordings were conducted in GFP-positive CINs. We found that the spontaneous firing frequency of DMS CINs was significantly lower in cocaine-administered mice than in saline controls (Fig. 1h; *t*_(19)_ = 2.30, *p* < 0.05). Since reversal training (Fig 1e, 1f) was performed approximately 24 d after the last cocaine injection, we also measured DMS CIN activity at this time point. We discovered that the spontaneous firing frequency was still lower in cocaine-administered mice than in saline controls (Fig. 1I; *t*_(19)_ = 2.42, *p* < 0.05), suggesting that cocaine induces long-lasting (24 d) inhibition of CIN activity.

To model voluntary cocaine-seeking behavior, we trained mice for intravenous self-administration (IVSA) of cocaine (Fig. 1j; Supplementary Fig. 1b), which is considered to be contingent drug exposure (Chen et al., 2008; Pascoli *et al*., 2014). The control group was yoked to receive saline infusions. Ten days after the last IVSA session, we found that the spontaneous firing frequency of DMS CINs was significantly lower in the cocaine-IVSA group than in the yoked saline controls (Fig. 1j; *t*_(24)_ = 3.31, *p* < 0.01). Taken together, these results suggest that experimenter-delivered (non-contingent i.p.) and voluntary cocaine administration (contingent IVSA) cause long-lasting inhibition of DMS CINs.

### Cocaine or alcohol exposure enhances the inhibition of CINs

Cocaine-mediated suppression of DMS CIN activity could result from an increase in inhibitory synaptic inputs (Gonzales *et al*., 2013; Guo *et al*., 2015). Therefore, we next tested whether repeated cocaine injections altered GABAergic inputs onto DMS CINs. ChAT-eGFP mice were i.p. injected with cocaine or saline for 7 d (Fig. 2a). Spontaneous inhibitory postsynaptic currents (sIPSCs) in DMS CINs were recorded 2 d after the last cocaine injection. We found that the frequency, but not the amplitude, of sIPSCs was greater in cocaine-exposed mice than in saline-injected controls (frequency: Fig. 2b, 2c, *t*_(28)_ = −2.23, *p* < 0.05; amplitude: Fig. 2d, *t*_(26)_ = 0.85, *p* > 0.05). In addition, we tested whether voluntary cocaine administration altered GABAergic transmission in DMS CINs. In mice trained for IVSA as described above (Fig. 1j, 2e), we discovered that cocaine IVSA also potentiated the sIPSC frequency, but not amplitude, in DMS CINs (frequency: Fig. 2f, *t*_(36)_ = −2.94, *p* < 0.01; amplitude: Fig. 2g, *t*_(36)_ = −1.68, *p* > 0.05). Moreover, we measured electrically evoked inhibitory postsynaptic currents (eIPSCs) in DMS CINs, and found that the eIPSC amplitude was significantly greater in the cocaine group than the saline-yoked controls (Fig. 2h; *F*_(1,16)_ = 18.02, *p* < 0.001). The paired-pulse ratio of the eIPSCs was significantly lower in the cocaine-IVSA group than in controls (Fig. 2i; *t*_(16)_ = 3.62, *p* < 0.01), indicating that cocaine IVSA increased the probability of presynaptic GABA release onto DMS CINs (Zucker and Regehr, 2002). Taken together, these results suggest that experimenter-delivered (non-contingent i.p.) and voluntary cocaine administration (contingent IVSA) potentiate GABAergic inputs onto DMS CINs.

**Figure 2.**
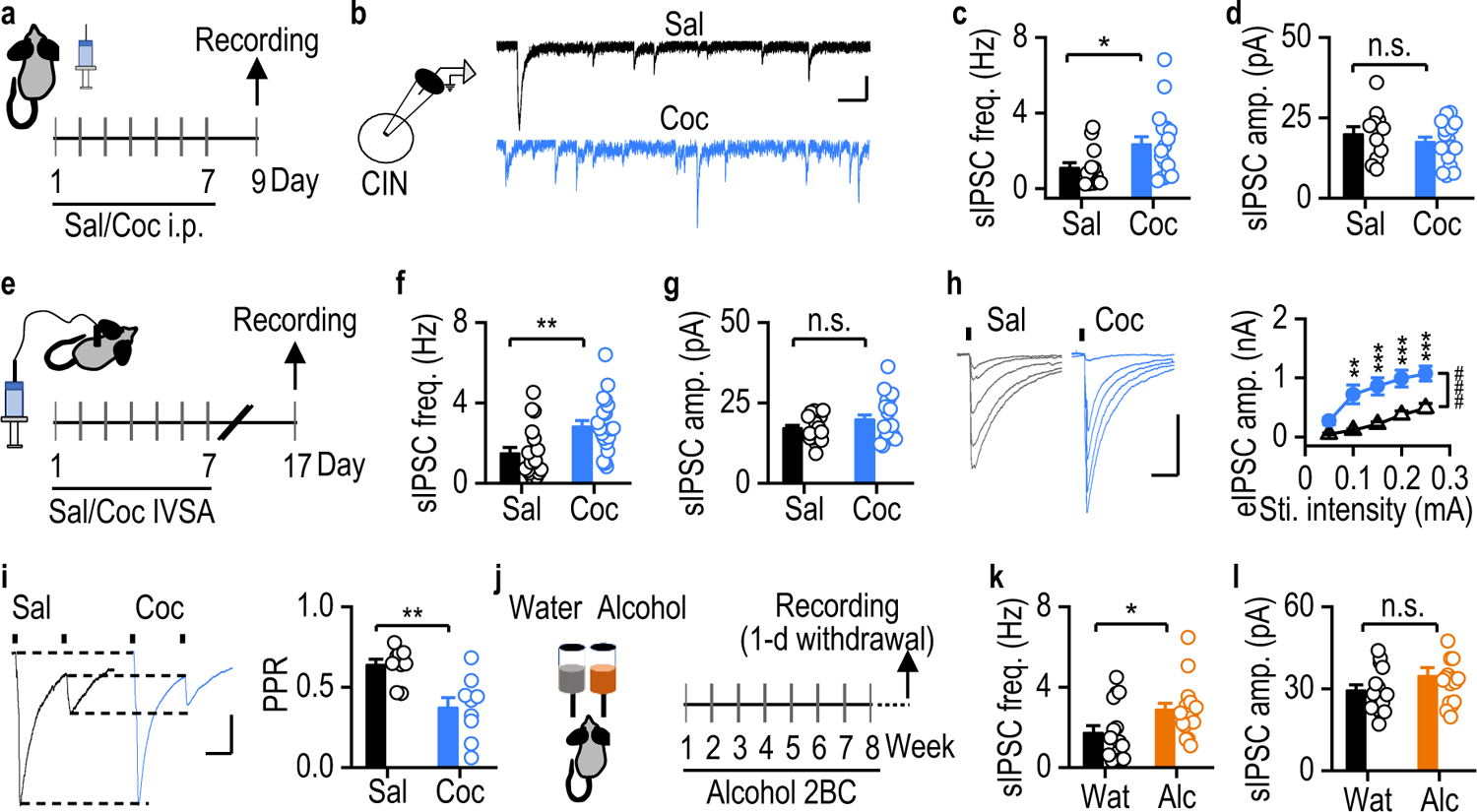
Cocaine or alcohol exposure potentiates GABAergic inputs onto DMS CINs. **a**, ChAT-eGFP mice were injected with saline or cocaine for 7 d and DMS CINs were recorded 2 d after the last injection. **b**, Traces of spontaneous IPSCs (sIPSCs). **c-d**, Cocaine increased sIPSC frequency (c) but not amplitude (d); \**p* < 0.05, n = 18/4 (Sal) and 20/4 (Coc). **e,** Mice were trained for cocaine IVSA for 7 d; CINs were recorded 10 d after the last session. **f-g**, Cocaine IVSA increased the frequency (f) but not amplitude (g) of sIPSCs; \*\**p* < 0.01, n = 13/4 (Sal) and 17/4 (Coc). **h**, Cocaine IVSA potentiated electrically evoked IPSCs (eIPSCs); ^###^*p* < 0.001 for the indicated comparison, two-way RM ANOVA; \*\*\**p* < 0.001, \*\**p* < 0.01 versus the saline group at the same stimulating intensity by *post-hoc* Tukey test; n = 9/4 (Sal) and 9/4 (Coc). **i**, Cocaine IVSA decreased the eIPSC paired-pulse ratio (PPR); \*\**p* < 0.01, n = 9/4 (Sal) and 9/4 (Coc). **j**, Mice were trained to self-administer water (Wat) or alcohol (Alc) for 8 weeks using the 2BC procedure. DMS CINs were recorded 1 d after the last alcohol session. **k-l**, Chronic alcohol consumption increased sIPSC frequency (k) but not amplitude (l); \**p* < 0.05, n = 15/3 (Wat) and 16/4 (Alc). Scale bars: 200 ms, 20 pA (b); 100 ms, 400 pA (h); 100 ms, 50 pA (i). Unpaired *t* test (c, d, f, g, i, k, l).

Since many substances of abuse cause reinforcement behaviors via similar neuronal mechanisms (Luscher, 2016; Pascoli *et al*., 2015), we also tested whether another widely used addictive substance, alcohol, altered GABAergic transmission onto DMS CINs. We trained ChAT-eGFP mice to self-administer alcohol in an intermittent-access two-bottle choice (2BC) procedure (Cheng *et al*., 2017; Hwa et al., 2011; Ma et al., 2018; Simms et al., 2008) (Fig. 2j). Again, we discovered that the frequency, but not the amplitude, of sIPSCs in DMS CINs was higher in alcohol-drinking mice than in water controls (frequency: Fig. 2k, *t*_(29)_ = −2.28, *p* < 0.05; amplitude: Fig. 2l, *t*_(29)_ = −1.32, *p* > 0.05). These results suggest that alcohol self-administration also potentiates GABAergic inputs onto DMS CINs.

### CINs mainly receive striatal inputs from dMSNs

Having shown that repeated cocaine or alcohol exposure potentiated presynaptic GABAergic inputs onto DMS CINs, we next examined the source of these inputs. A previous study found that the majority of inputs to CINs originated from the striatum itself (Guo *et al*., 2015). The striatum comprises two equal populations of principal MSNs that express either dopamine D1 receptors (D1Rs) or D2Rs (Cheng *et al*., 2017; Gerfen and Surmeier, 2011; Kreitzer and Malenka, 2008). D1R-MSNs and D2R-MSNs are located in the direct- and indirect-pathways of the basal ganglia and thus are also referred to as dMSNs or iMSNs, respectively (Cheng *et al*., 2017; Gerfen and Surmeier, 2011; Kreitzer and Malenka, 2008).

To examine whether CINs received equal or distinct afferent inputs from dMSNs and iMSNs, we used rabies-mediated monosynaptic retrograde tracing (Wickersham et al., 2007). The helper viruses, AAV-DIO-TVA-mCherry and AAV-DIO-RG, were infused into the DMS of ChAT-Cre;D1tdT mice, where CINs expressed Cre recombinase and dMSNs expressed tdTomato (tdT; red) (Fig. 3a). This infusion caused selective expression of TVA-mCherry and RG in CINs. Three weeks later, rabies-GFP was infused at the same location in the DMS (Fig. 3a right, 3b), leading to selective rabies-GFP infection of TVA-expressing CINs. RG allowed the transfer of rabies-GFP from CINs to their presynaptic neurons (Fig. 3c). These CINs, referred to as “starter CINs”, expressed mCherry and GFP and thus appeared yellow. Similarly, dMSNs presynaptic to the starter CINs contained tdT and GFP and were also yellow (Fig. 3d-3h). To differentiate starter CINs from presynaptic dMSNs, we performed ChAT immunostaining to label all CINs using Alexa Fluor 647, which was far-red (pseudocolored with cyan) (Fig. 3g, 3h). Consequently, the starter CINs were labeled with mCherry, GFP, and cyan, whereas presynaptic dMSNs expressed tdT and GFP. In addition, striatal neurons that expressed only GFP were presynaptic non-dMSNs. Since over 90% of striatal neurons are either dMSNs or iMSNs (Gerfen and Surmeier, 2011; Kreitzer and Malenka, 2008), we assumed that most of these “GFP only” neurons were putative presynaptic iMSNs. We discovered a higher number of presynaptic dMSNs than putative presynaptic iMSNs (Fig. 3i; *t*_(68)_ = 7.87, *p* < 0.001). Additionally, we found a few CINs (co-labeled with GFP and cyan) that formed monosynaptic connections with the starter CINs (Dorst et al., 2020; Sullivan et al., 2008). These findings suggest that CINs receive striatal inputs that mainly originate from dMSNs.

**Figure 3.**
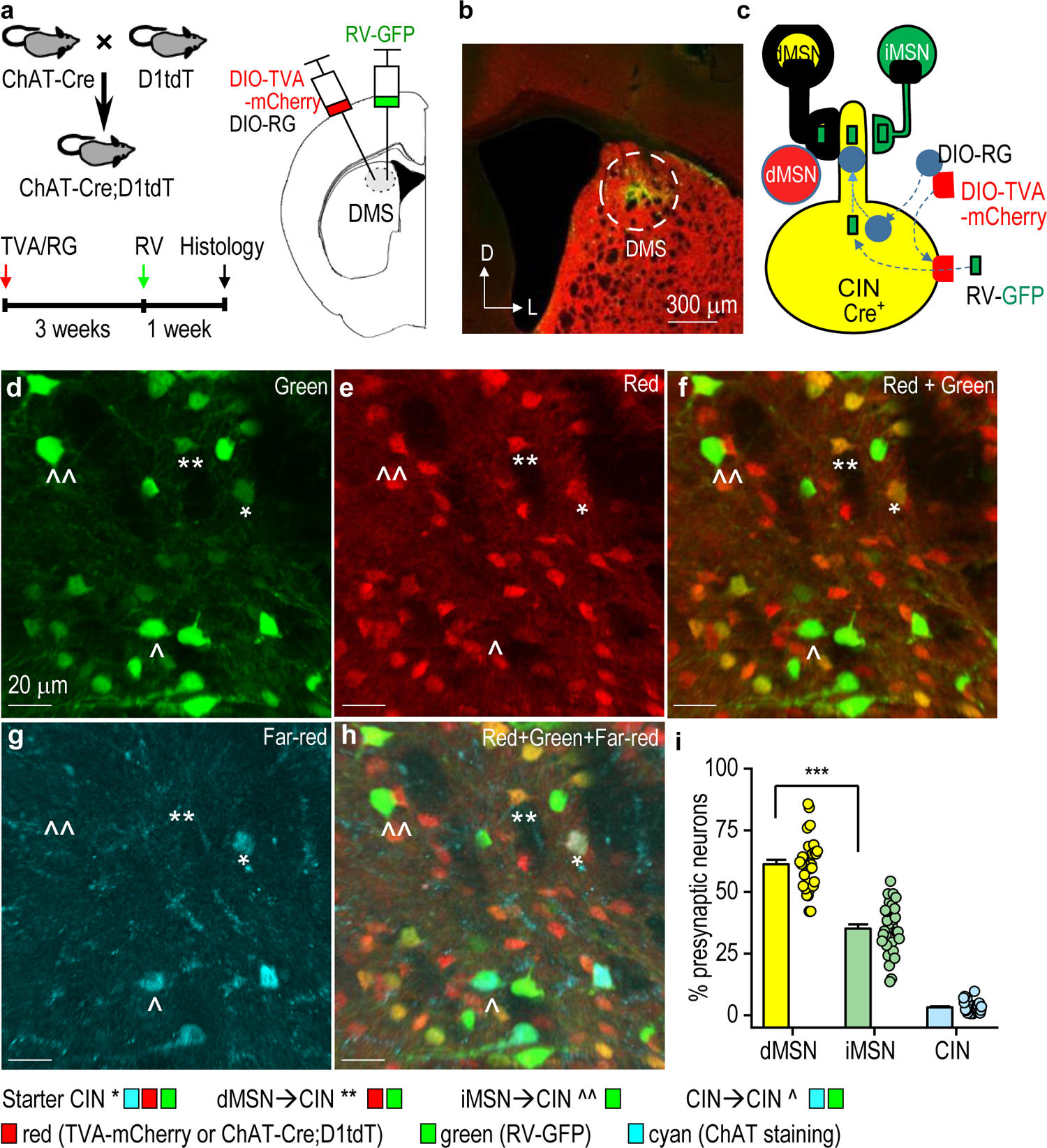
CINs primarily receive striatal inputs from dMSNs. **a**, Schematic of the experimental design. Cre-dependent helper viruses (AAV-DIO-TVA-mCherry and AAV-DIO-RG) were infused into the DMS of ChAT-Cre;D1tdT mice, followed by rabies virus (RV-GFP) infusion at the same site 3 weeks later. Coronal sections were prepared 1 week after rabies virus infusion and were stained for choline acetyltransferase (ChAT; far-red, pseudocolored with cyan). **b,** Confocal micrograph showing the injection site and viral expression in the DMS. **c**, Schematic showing viral expression and retrograde spread of RV-GFP. AAV-DIO-TVA-mCherry and AAV-DIO-RG infected Cre-positive CINs. RV-GFP infected TVA-positive CINs (starter cells expressed GFP and mCherry), labeling their presynaptic neurons with GFP. Since dMSNs were labeled red (from D1tdT), dMSNs presynaptic to the starter CINs were yellow (red and green overlap), whereas putative iMSNs were green. **d-h**, Confocal micrographs of a DMS section. **i**, Summarized data showing that CINs received significantly more inputs from dMSNs than from iMSNs; \*\*\**p* < 0.001 by unpaired *t* test, n = 35 slices from 4 mice. RV: rabies virus; RG: rabies glycoprotein; TVA, EnvA receptor.

### CINs receive stronger inhibition from dMSNs than from iMSNs

Since MSNs are GABAergic,the greater number of anatomic inputs from dMSNs than from iMSNs indicates that CINs are primarily inhibited by dMSNs. To test this, we generated D1-Cre;Ai32 and A2a-Cre;Ai32 mice, in which dMSNs and iMSNs, respectively, expressed ChR2 to allow selectively optogenetic stimulation (Fig. 4a, left). CINs in DMS slices were identified by their large size, spontaneous firing, and characteristic sag at hyperpolarizing currents (Fig. 4a, right). Optical dMSN excitation induced a large inhibitory postsynaptic current (oIPSC) in CINs that was blocked by the GABA_A_R antagonist, picrotoxin (Fig. 4b). Importantly, we discovered that dMSN→CIN oIPSCs were larger than iMSN→CIN oIPSCs (Fig. 4c; *F*_(1,8)_ = 16.83, *p* < 0.01). To further confirm this finding, we generated another two lines of transgenic mice that expressed a different excitatory opsin, Chrimson, in either dMSNs or iMSNs. Moreover, CINs were directly identified by their expression of GFP in these animals. Again, we found greater dMSN→CIN oIPSCs than iMSN→CIN oIPSCs (Fig. 4d; *t*_(24)_ = 3.95, *p* < 0.001). Notably, these transgenic mice expressed the opsins in D1R- and A2AR-containing neurons throughout their brains (Lu et al., 2021a; Wei et al., 2018). Next, we virally expressed Chrimson in DMS dMSNs or iMSNs in D1-Cre;ChATeGFP and A2A-Cre;ChATeGFP mice (Fig. 4e, 4f). Whole-cell recordings revealed greater dMSN→CIN oIPSCs than iMSN→CIN oIPSCs. (Fig. 4g; *t*_(22)_ = 3.38, *p* < 0.01). These results suggest that CINs receive stronger inhibitory inputs from dMSNs than from iMSNs.

**Figure 4.**
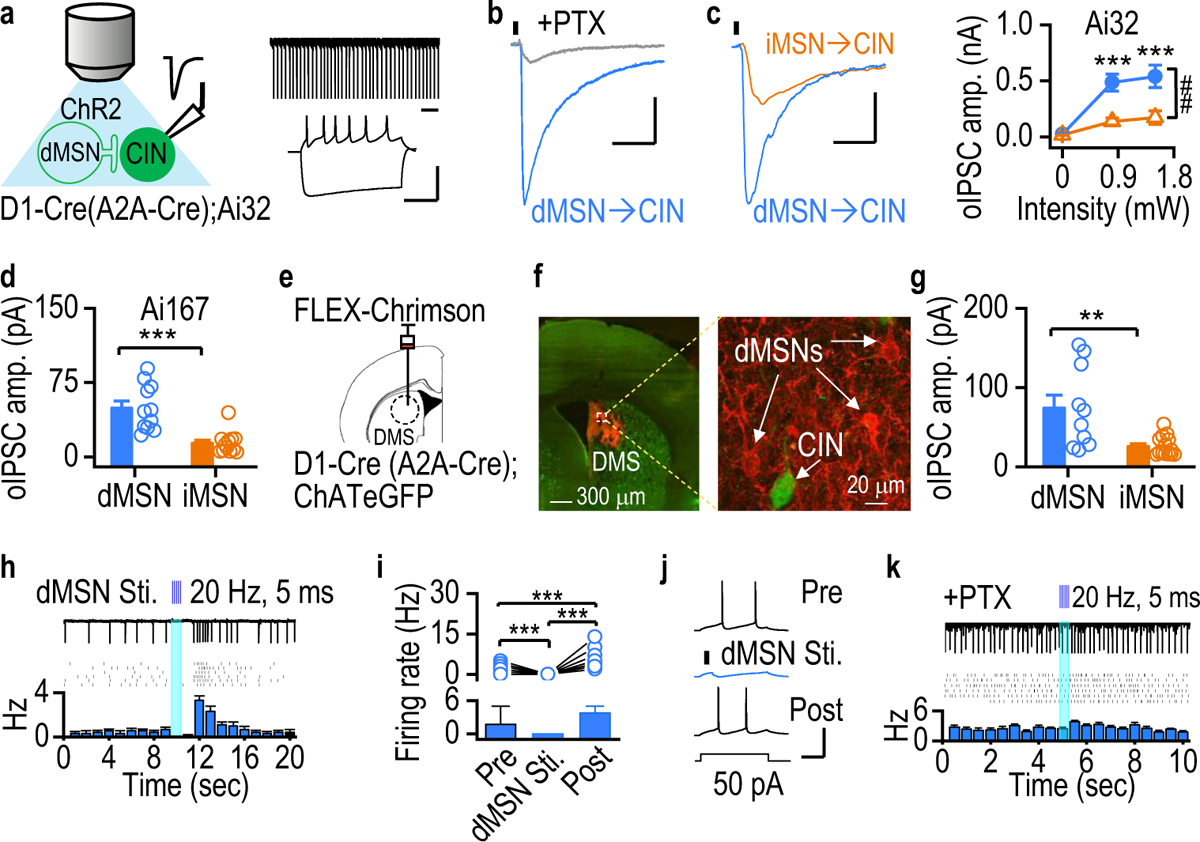
CINs receive stronger functional GABAergic inputs from dMSNs than from iMSNs. **a**, Optogenetic stimulation of ChR2-expressing dMSNs (left) and physiological identification of DMS CINs (right) in D1-Cre;Ai32 and A2A-Cre;Ai32 mice, which selectively expressed ChR2 in dMSNs and iMSNs, respectively. Trace of CIN recording showing spontaneous firing (top right) and characteristic sag (bottom right). **b**, Picrotoxin (PTX, 100 µM) blocked the optically evoked IPSCs from dMSNs to a CIN (oIPSC^dMSN→CIN^). **c**, oIPSC^dMSN→CIN^ (n = 4/3) was greater than oIPSC^iMSN→CIN^ (n = 6/2); ^##^*p* < 0.01 for the indicated comparison, two-way RM ANOVA; \*\*\**p* < 0.001 versus the iMSN→CIN group at the same intensity, Tukey *post-hoc* test. **d**, oIPSC^dMSN→CIN^ (n = 14/7) was greater than oIPSC^iMSN→CIN^ (n = 12/2) using Ai167 mice that Cre-dependently expressed Chrimson-tdT in dMSNs (D1-Cre;Ai167;ChATeGFP mice) and in iMSNs (A2A-Cre;Ai167;ChATeGFP mice); \*\*\**p* < 0.001. **e**, AAV-FLEX-Chrimson-tdT was infused into the DMS of D1-Cre (A2A-Cre);ChATeGFP. **f**, Image of Chrimson-tdT expression in dMSNs and GFP in CINs. **g**, oIPSC^dMSN→CIN^ (n = 10/3) was greater than oIPSC^iMSN→CINs^ (n = 14/5) in mice that selectively expressed Chrimson-tdT in the DMS; \*\**p* < 0.01. **h**, Recording of CIN pause-rebound firing after a burst of dMSN stimulation in a D1-Cre;Ai32 mouse. Top, sample firing; middle, multiple sweeps; bottom, the corresponding peristimulus histogram. **i**, Summarized data demonstrating pause-rebound CIN firing after dMSN stimulation; \*\*\**p* < 0.001. one-way RM ANOVA followed by Tukey *post-hoc* test, n = 44 sweeps from 8 neurons. **j**, dMSN stimulation (50 Hz, 2 pulses) reversibly inhibited evoked CIN firing. **k**, Picrotoxin abolished the CIN pause-rebound response after a burst of dMSN stimulation. Scale bars: 1 sec (a, top right); 100 ms, 50 mV (a, bottom right); 50 ms, 200 pA (b); 15 ms, 200 pA (c); 100 ms, 50 mV (j). Unpaired *t* test (d, g).

To investigate the consequences of dMSN-mediated inhibition of CINs, we measured spontaneous CIN firing following a burst of light stimulation of dMSNs in DMS slices from D1-Cre;Ai32 mice. We observed a strong pause followed by a rebound in CIN firing (Supplementary Fig. 2; Fig. 4h, 4i; *F*_(2,129)_ = 68.03, *p* < 0.001). We also discovered that optical dMSN stimulation inhibited the CIN firing evoked by current injection (Fig. 4j). Additionally, picrotoxin abolished the inhibitory effect of dMSN stimulation on CIN firing activity (Fig. 4k). These results suggest that dMSN excitation is sufficient to suppress CIN firing activity.

### Cocaine and alcohol potentiate inhibitory dMSN**→**CIN transmission

A large body of literature shows that chronic exposure to addictive substances, including cocaine or alcohol, potentiates dMSN activity (Cheng *et al*., 2017; Dong et al., 2017; Luscher, 2016; Ma *et al*., 2018; Roberts-Wolfe et al., 2018). Next, we tested whether administration of cocaine or alcohol also potentiated the inhibitory dMSN→CIN transmission. While cocaine exposure has been known to potentiate glutamatergic inputs onto dMSNs in the ventral striatum (Kauer and Malenka, 2007; Ungless et al., 2001), it was not known whether this potentiation also occurred in the DMS. To test this, mice were injected with cocaine for 7 d, and DMS dMSNs were recorded 2 d after the last cocaine injection (Fig. 5a). We found that the AMPAR/NMDAR ratio in DMS dMSNs was significantly higher in cocaine-treated mice than in saline controls (Fig. 5b; *t*_(15)_ = −3.03, *p* < 0.01). Additionally, AMPA-induced currents in DMS dMSNs were greater in cocaine-administered mice than in saline controls (Fig. 5c; *t*_(15)_ = −3.03, *p* < 0.01). To test whether this cocaine-mediated potentiation enhanced downstream dMSN→CIN transmission, we recorded DMS CINs in the same cocaine-injected mice (Fig. 5a). dMSN→CIN oIPSCs were measured 2 d and 24 d after the last injection. We discovered that the oIPSCs were significantly higher in cocaine-administered mice than in saline controls at both time points (2 d: Fig. 5d, *F*_(1,45)_ = 0.74, *p* < 0.05; 24 d: Fig. 5e, *t*_(22)_ = −2.22, *p* < 0.05). These results suggest that experimenter-delivered (noncontingent i.p.) cocaine potentiates glutamatergic transmission in dMSNs and causes a long-lasting potentiation of inhibitory dMSN→CIN transmission in the DMS.

**Figure 5.**
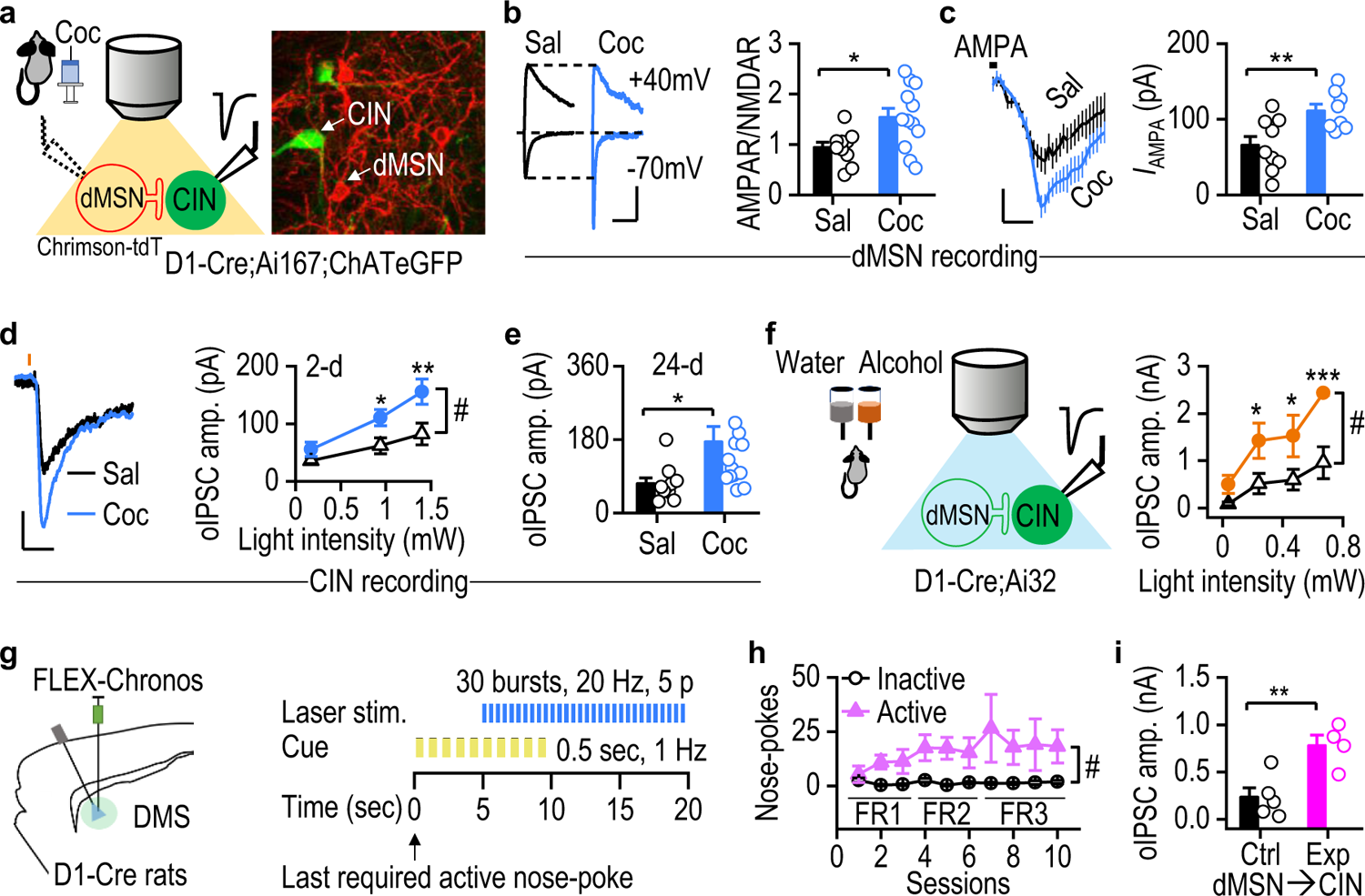
Chronic cocaine or alcohol administration potentiates GABAergic dMSN→CIN transmission in the DMS. **a, b**, Repeated cocaine injections increased AMPAR/NMDAR ratios in dMSNs of D1-Cre;Ai167;ChATeGFP mice. The animals expressed Chrimson-tdT in dMSNs and eGFP in CINs (a). EPSCs were electrically evoked (b); \**p* < 0.05, n = 10/3 (Sal) and 12/2 (Coc). **c**, Cocaine potentiated the AMPA (5 μM)-induced current (*I*_AMPA_) in dMSNs; \*\**p* < 0.01, n = 9/3 (Sal) and 8/3 (Coc). **d**, **e,** Cocaine potentiated oIPSC^dMSN→CIN^ 2 d (d) and 24 d (e) after the last injection; \*\**p* < 0.01, \**p* < 0.05, n = 22/5 (Sal, d) and 25/7 (Coc, d), 10/2 (Sal, e) and 14/4 (Coc, e). **f,** Alcohol chronically potentiated oIPSC^dMSN→CIN^. Mice were trained to consume alcohol using the 2BC procedure (left) and oIPSCs were recorded 1 d after the last drinking session (right); ^#^*p* < 0.05 for the indicated comparison; \*\*\**p* < 0.001, \**p* < 0.05 versus water at the same intensity, n = 6/4 (Water) and 5/3 (Alcohol). **g**, AAV-FLEX-Chronos-GFP was infused into the DMS of D1-Cre rats (left). Animals were trained to nose-poke for optical stimulation of dMSNs (30 bursts, separated by 250 ms) (right). **h**, Animals learned to nose-poke the active port to receive dMSN stimulation; ^#^*p* < 0.05 for the indicated comparison**. i**, dMSN self-stimulation potentiated oIPSC^dMSN→CIN^ 5 d after the last training session; \*\**p* < 0.01, n = 5/5 (Ctrl) and 4/1 (Exp). Scale bars: 100 ms, 50 pA (b); 2 min, 30 pA (c); 100 ms, 20 pA (d). Unpaired *t* test (b, c, e, i); two-way RM ANOVA followed by Tukey *post-hoc* test (d, f, h).

We also tested whether chronic exposure to another addictive substance, alcohol, altered dMSN→CIN transmission. Mice were trained to consume alcohol for 8 weeks using the intermittent-access 2BC procedure (Fig. 5f). Control mice only had access to water. One day after the last alcohol session, we prepared DMS slices and recorded dMSN→CIN oIPSCs. The oIPSCs were significantly greater in alcohol-drinking mice than in controls (Fig. 5f; *F*_(1,9)_ = 10.33, *p* < 0.05). This result suggests that chronic alcohol consumption potentiates inhibitory dMSN→CIN transmission in the DMS.

Our results suggest that dMSN potentiation by cocaine or alcohol exposure enhances inhibitory dMSN→CIN transmission. Similarly, dMSN activity can be potentiated by natural reward self-administration (Shan *et al*., 2014) or intracranial self-stimulation of either dMSNs or dopaminergic neurons (Lalive *et al*., 2018; Pascoli *et al*., 2015). Therefore, we next examined whether intracranial self-stimulation of DMS dMSNs also potentiated dMSN→CIN transmission. We virally expressed Chronos in DMS dMSNs of D1-Cre rats and then trained them to nose-poke to receive dMSN stimulation (Fig. 5g, 5h). We found that some rats exhibited behavioral reinforcement for dMSN excitation, while others did not (Fig. 5h; *F*_(1,4)_ = 8.32, *p* < 0.05). Five days after the last training session, we prepared DMS slices and recorded dMSN→CIN oIPSCs. We observed that these oIPSCs were stronger in rats that had exhibited self-stimulation than in those that did not (Fig. 5I; *t*_(7)_ = 3.61, *p* < 0.01, Supplementary Fig 3). These data suggest that intracranial self-stimulation of dMSNs also potentiates inhibitory dMSN→CIN transmission in the DMS.

Taken together, these findings indicate that both reinforcement of addictive substances such as cocaine and alcohol and dMSN self-stimulation potentiates inhibitory dMSN→CIN transmission.

### Chemogenetic CIN inhibition impairs reversal learning

The studies described above indicate that repeated exposure to addictive substances enhances inhibitory dMSN→CIN transmission and thus suppresses CIN activity in the DMS. Because CINs are known to regulate reversal learning (Aoki *et al*., 2015; Bradfield *et al*., 2013; Galaj *et al*., 2019; Matamales *et al*., 2016), we next tested whether chemogenetic inhibition of DMS CINs also altered cognitive flexibility. An inhibitory DREADD, hM4Di, was selectively expressed in DMS CINs of ChAT-Cre mice (Fig. 6a, 6b). These animals were then trained on the two action-outcome reversal learning task described in Figure 1 (Fig. 6c). We observed that both hM4Di-infused and control groups preferred to nose-poke for food over sucrose (Supplementary Fig. 4a; *F*_(1,16)_ = 6.90, *p* < 0.05) and we therefore analyzed food-seeking and -taking behaviors in subsequent studies. Nose-pokes or magazine entries did not differ between groups across sessions during initial learning (Fig. 6d, 6e). After initial learning, the action-outcome contingencies of the operant task were reversed (Fig. 6f). Thirty minutes before each reversal training session, mice were administered with clozapine N-oxide (CNO) to suppress CIN activity via hM4Di. While nose-pokes did not differ between groups, the hM4Di group showed more total magazine entries on the first day of reversal learning than the controls (nose-pokes: Fig. 6g, *F*_(1,17)_ = 0.87, *p* > 0.05; magazine entries: Fig. 6h, *F*_(1,17)_ = 0.18, *p* < 0.05). These data suggest that CIN activity is required for reward-taking (i.e., magazine entry) but not for reward-seeking (i.e., nose-poking) behavior during reversal learning.

**Figure 6.**
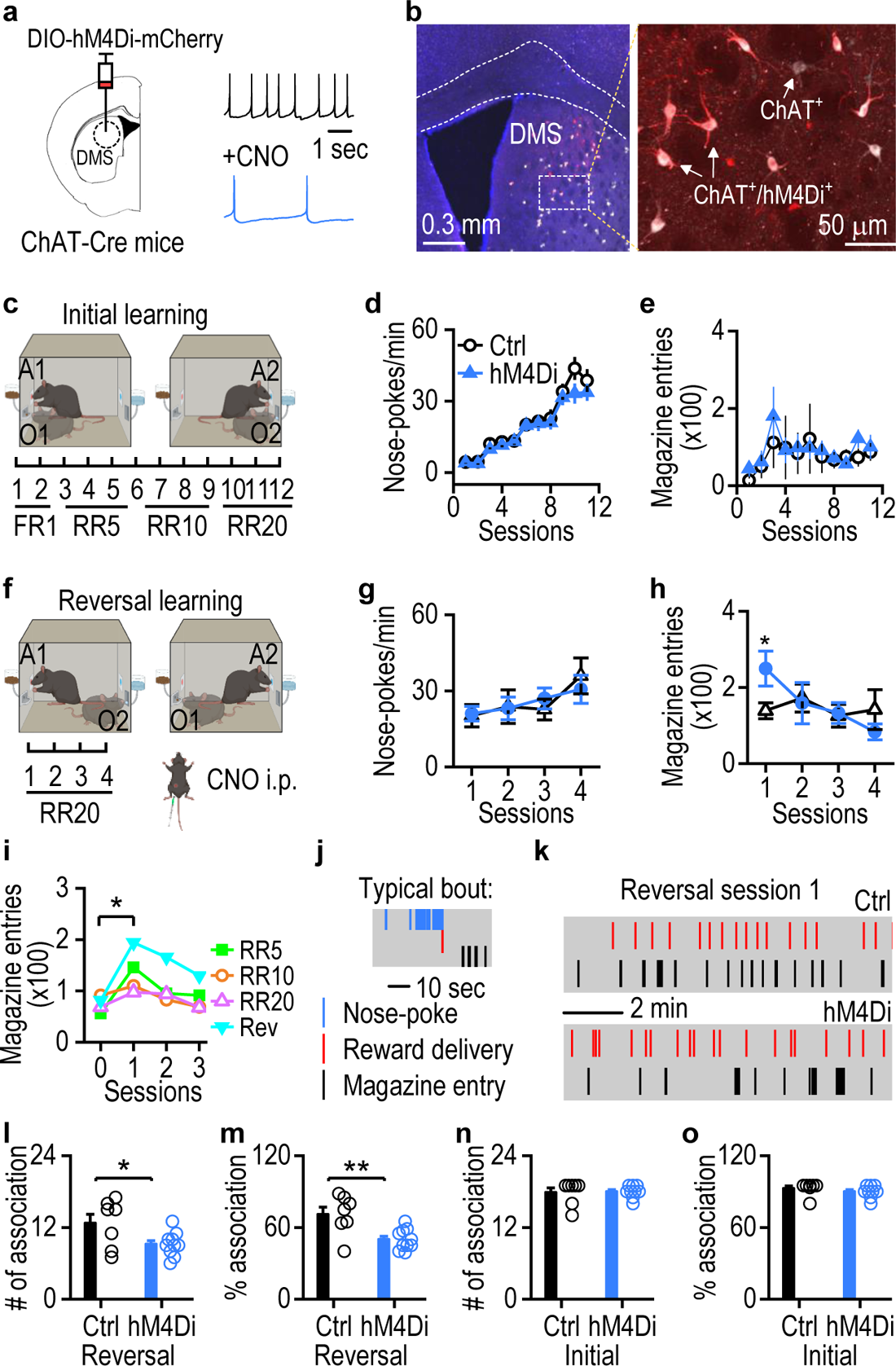
Chemogenetic inhibition of DMS CINs during reversal learning impairs action-outcome association. **a**, AAV-DIO-hM4Di-mCherry was infused into the DMS of ChAT-Cre mice. **b**, Image of hM4Di-mCherry expression in CINs, with ChAT-staining (white). **c**, Initial action-outcome contingency training (A1→O1; A2→O2). **d**, **e,** Nose-poking for food (d) and magazine entries (e) did not differ between groups during initial learning; n = 10 (hM4Di) and 9 (Ctrl, mCherry) mice. **f**, Reversal learning (A1→O2; A2→O1). CNO (3-5 mg/kg) was administered 30 min before each session. **g**, Nose-poking for food did not differ between groups during reversal learning. **h**, Chemogenetic CIN inhibition via hM4Di increased magazine entries for food in the first reversal session; \**p* < 0.05 versus controls for the same session, Tukey *post hoc* test. **i**, Food magazine entries were higher on day one of a new schedule (session 1) than on the last day of the previous schedule (session 0); \**p* < 0.05, n = 20 mice. **j**, A representative action-outcome association bout. **k**, Example showing decreased magazine entries for food collection after CIN inhibition on reversal day one. **l**, **m**, Bouts (l) and percentage bouts (m) with reward delivery and successful collection were lower in the hM4Di group than in controls on reversal day one; \**p* < 0.05, \*\**p* < 0.01. **n**, **o**, Bouts (n) and percentage bouts (o) did not differ between groups on the last day of initial learning; *p* > 0.05. Unpaired *t* test: (i, l-o); two-way RM ANOVA (d, e, g, h).

This finding is consistent with the fact that the animals detect a change in the action-outcome contingency during reversal training and that CINs are known to respond to salient environmental changes (Morris et al., 2004). If CIN activity is inhibited, the animals may be less likely to detect the change in outcome, leading to repeated magazine entries. We propose that upon detection of an altered action-outcome contingency, animals increase magazine entries to confirm this change. To investigate this, we analyzed magazine entries during initial learning. This analysis supported our hypothesis because we discovered that magazine entry increased when the training schedule of the task changed, and then declined with continued training (Fig 6i; *t*_(6)_ = −3.03, *p* < 0.05).

Next, we analyzed the first day of reversal training for both groups in greater detail (Fig. 6j). We found that while the control mice entered the magazine port to collect rewards immediately after food delivery, hM4Di mice often did not enter the port after each delivery (Fig. 6k). Consequently, we found that the number of successful bouts of reward delivery and collection, as well as the percentage of successful bouts, were significantly lower in the hM4Di group than in the control group (number: Fig. 6l, *t*_(15)_ = − 2.40, *p* < 0.05; percentage: Fig. 6m, *t*_(15)_ = −3.42, *p* < 0.01). This result contrasted with the successful bouts on the last day of initial training, during which there was no difference between the groups (number: Fig. 6n, *t*_(19)_ = 0.40, *p* > 0.05; percentage: Fig. 6o, *t*_(19)_ = − 0.33, *p* > 0.05). Together, these results suggest that DMS CINs are required for at least the early stage of reversal learning.

### Time-locked optogenetic CIN inhibition impairs reversal and extinction learning

Having shown that DMS CINs are required to reverse instrumental learning, the time point at which this occurs was still unclear, as CNO-mediated CIN inhibition lasts for hours. CINs are activated by salient environmental stimuli (Morris *et al*., 2004), including changes in reward delivery (e.g., food→sucrose) during the reversal task. Therefore, we hypothesized that DMS CIN activity during reward delivery would regulate reversal learning. To test this hypothesis, we used optogenetics to selectively inhibit DMS CINs while rewards were delivered during reversal training sessions in rats that are known to demonstrate strong goal-directed learning (Bradfield *et al*., 2013). An inhibitory opsin, eNpHR, was virally expressed in DMS CINs of ChAT-Cre;tdT rats (Fig. 7a, 7b). These animals were then trained on the two action-outcome reversal learning task (Fig. 7c). During initial learning, the control and eNpHR groups did not differ in lever pressing or magazine entries (lever pressing: Supplementary Fig. 5a, *F*_(1,15)_ = 0.61, *p* > 0.05; entries: Supplementary Fig. 5b, *F*_(1,15)_ = 0.09, *p* > 0.05). Both groups also demonstrated significant devaluation (Fig 7d; control: Fig. 7e left, *t*_(14)_ = 3.50, *p* < 0.01; eNpHR: Fig. 7e right, *t*_(14)_ = 2.82, *p* < 0.05) and had similar devaluation indices (Fig. 7f; *t*_(14)_ = 0.73, *p* > 0.05).

**Figure 7.**
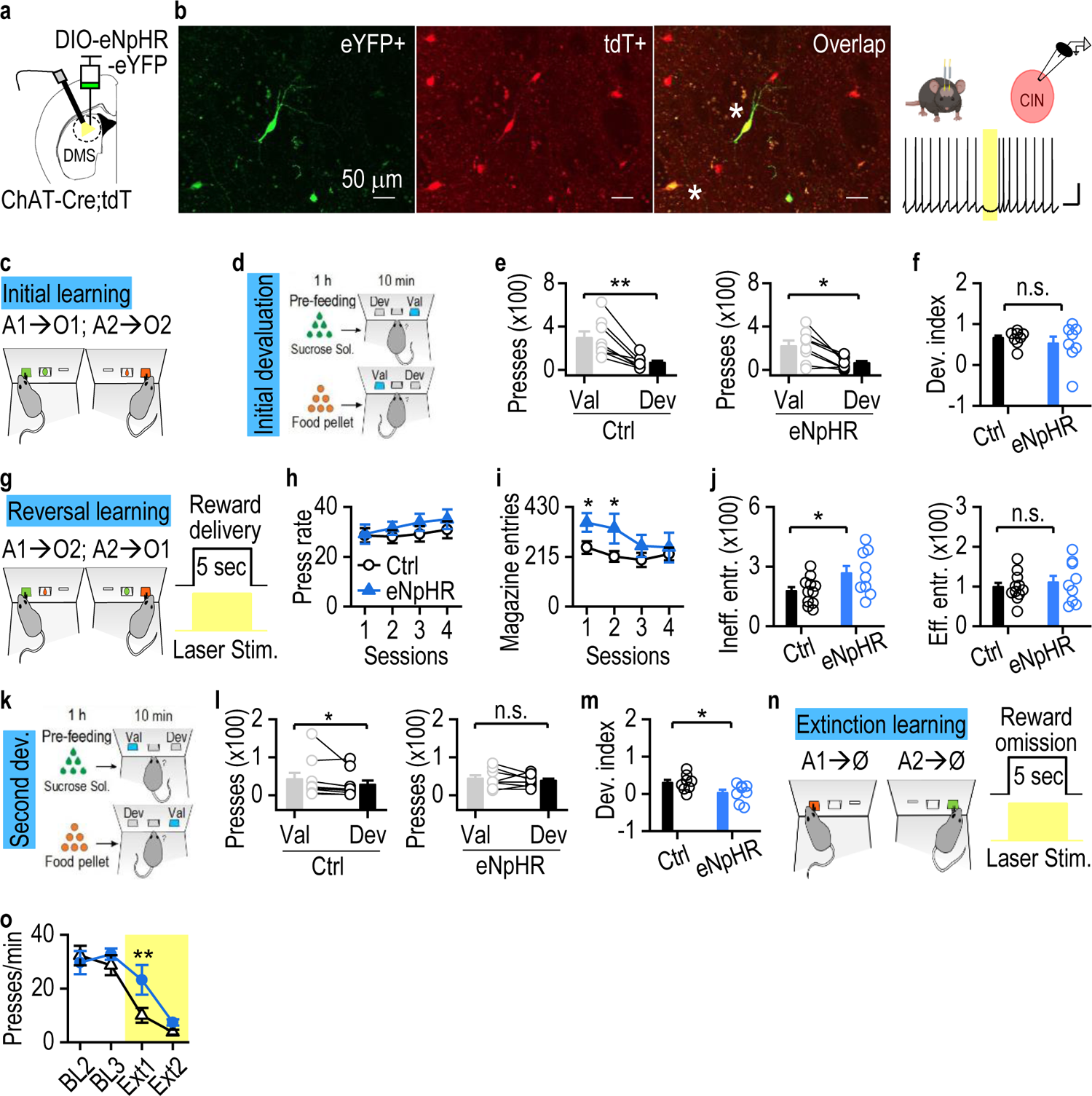
Time-locked optogenetic inhibition of DMS CINs impairs reversal and extinction learning. **a,** AAV-DIO-eNpHR-eYFP was infused into the DMS of ChAT-Cre;tdT rats. **b**, tdT-positive CINs expressed eYFP (left) and were inhibited by yellow light stimulation of eNpHR (right). **c**, Initial action-outcome contingency training (A1→O1; A2→O2). **d**, Initial devaluation test. Val: valued, Dev: devalued. **e,** Control (eYFP only) and eNpHR groups exhibited successful devaluation; \*\**p* < 0.01, \**p* < 0.05, n = 8 rats/group. **f**, Initial devaluation index did not differ between groups. **g**, Reversal training (A1→O2; A2→O1). Reward deliveries were paired with laser stimulation. **h**, Lever pressing did not differ between groups during reversal training. **i**, Optogenetic inhibition of DMS CINs increased magazine entries during the first two reversal sessions; \**p* < 0.05. **j**, Ineffective (left), but not effective (right), magazine entries were higher in the eNpHR group than in controls on reversal day one; \**p* < 0.05. **k**, Second devaluation test. **l**, The control group exhibited successful devaluation (left, \**p* < 0.05), whereas the eNpHR group did not (right). **m,** The devaluation index was lower in the eNpHR group than in controls; \**p* < 0.05. **n**, Schematic for extinction training. Reward omissions were paired with laser stimulation. **o**, Lever press rate was higher in the eNpHR group than in controls on extinction day one; \*\**p* < 0.01; n = 6 (Ctrl) and 7 (eNpHR) rats. Paired *t* test (e, l); unpaired *t* test (f, j, m); two-way RM ANOVA followed by Tukey *post-hoc* test (h, i, o). Scale bars: 1 sec, 20 mV (b).

The rats then underwent reversal training, during which constant light stimulation to inhibit DMS CINs was time-locked to the reward deliveries (Fig. 7g). We discovered that although lever pressing performance was unaffected (Fig. 7h; *F*_(1,15)_ = 0.95, *p* > 0.05), the eNpHR group made significantly more magazine entries than the controls during the first two sessions (Fig. 7i; *F*_(1,15)_ = 3.24, *p* < 0.05). We further classified magazine entries into two types: effective and ineffective, depending on whether the rat entered the magazine during a reward delivery or at another time, respectively. We found that the total number of ineffective entries on the first day of reversal training was significantly higher in the eNpHR group than in the control group (Fig. 7j left; *t*_(18)_ = −2.24, *p* < 0.05), whereas effective entries did not differ between groups (Fig. 7j right; *t*_(18)_ = −0.57, *p* > 0.05). Additionally, the return maps of inter-entry intervals (Matamales *et al*., 2020) revealed that the center of the data points was closer to the origin for the eNpHR group than for the controls (Supplementary Fig. 5c, 5d). These results demonstrate that the eNpHR rats checked the magazine for a reward more frequently than the control animals, although lever pressing performances did not differ between these two groups.

Following reversal training, we performed devaluation testing (Fig. 7k) and found that while the control group could be devalued, the eNpHR group could not (control: Fig 7l left, *t*_(7)_ = 2.72, *p* < 0.05; eNpHR: Fig 7l right, *t*_(7)_ = 0.82, *p* > 0.05). Consequently, the devaluation index was significantly lower in the eNpHR group than in controls (Fig. 7m; *t*_(14)_ = 2.34, *p* < 0.05), suggesting that inhibition of DMS CINs during reward delivery impaired reversal learning. Importantly, optogenetic CIN inhibition time-locked to lever presses did not affect reversal learning (Supplementary Fig. 5e, *t*_(9)_ = 0.72, *p* > 0.05).

Reversal learning involves the inhibition of established action-outcome contingencies (extinction) and the formation of new contingencies. Therefore, we next tested whether inhibition of DMS CINs also altered extinction learning (Fig. 7n). Optical stimulation was used to inhibit DMS CINs when a reward was supposed to be delivered. We observed a higher number of lever presses by the eNpHR group on the first day of extinction, as compared with the control group (Fig. o, *F*_(1,11)_ = 1.03, *p* < 0.05), suggesting that inhibition of DMS CINs delayed extinction learning. Taken together, these results indicate that time-locked inhibition of DMS CINs during reward delivery impairs reversal and extinction learning.

### SNr-projecting dMSNs send collaterals to inhibit CINs

So far, we have shown that reinforcing behaviors induced by cocaine, alcohol, or dMSN self-stimulation potentiate inhibitory dMSN→CIN transmission, and thereby reduce cognitive flexibility. However, the dMSN→SNr pathway has also been shown to mediate the reinforcing effects of cocaine and alcohol (Hellard *et al*., 2019). Additionally, animals have been shown to self-stimulate dMSNs and self-inhibit SNr neurons (Lalive *et al*., 2018). Therefore, we next asked whether SNr-projecting dMSNs that mediate reinforcement (Freeze et al., 2013; Lalive *et al*., 2018) also send collaterals to inhibit CIN activity.

To test this, we infused AAVretro-DIO-ChR2-eGFP into the SNr of D1-Cre;ChATeGFP mice (Fig. 8a). This infusion led to a retrograde expression of ChR2-eGFP on the cell membrane of dMSNs (Fig. 8b). Optical stimulation of ChR2 evoked a direct depolarization current in dMSNs (Supplementary Fig. 6) and an oIPSC in CINs that was blocked by picrotoxin (Fig. 8c). Additionally, a burst of light stimulation of dMSNs was sufficient to suppress CIN firing (Fig. 8d). These results suggest that SNr-projecting dMSNs send axonal collaterals to inhibit DMS CINs. Furthermore, we examined whether there were any axonal fibers of CIN-projecting dMSNs in the internal globus pallidus (GPi) and SNr regions of the rabies-GFP-infused ChAT-Cre;D1tdT mice described previously (Fig. 3, 8e). We discovered that the dMSNs (green and red) that innervated CINs also projected to the GPi (Fig. 8f-h) and SNr (Fig. 8i-k). Taken together, our results suggest that dMSNs send axons to the SNr to form a connection that mediates reinforcement (Hellard *et al*., 2019; Lalive *et al*., 2018), and also send collateral axons to CINs to inhibit cognitive flexibility.

**Figure 8.**
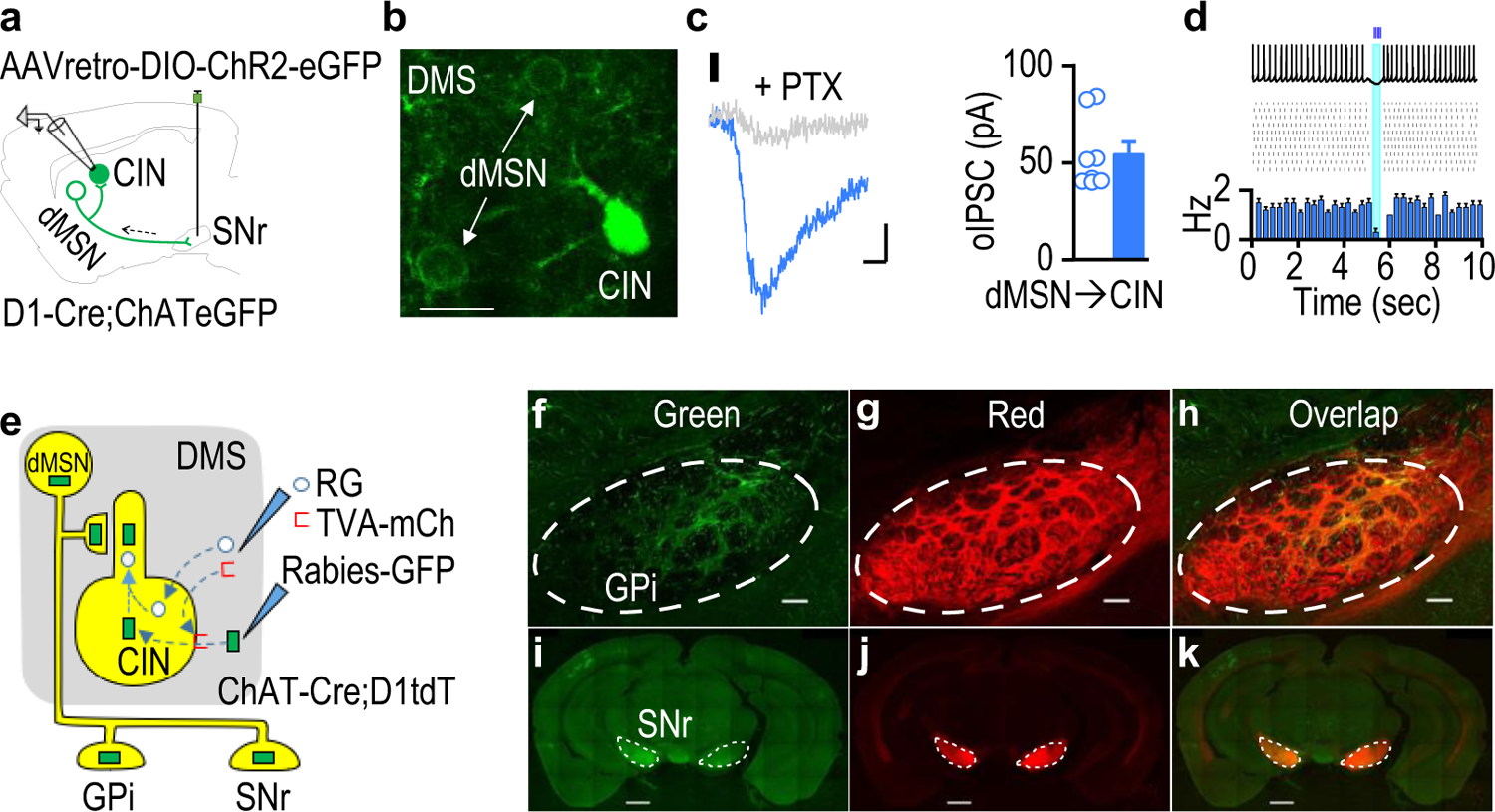
SNr-projecting dMSNs send collaterals to inhibit CINs in the DMS. **a**, AAVretro-DIO-ChR2-eGFP was infused into the SNr of D1-Cre;ChATeGFP mice and DMS CINs were recorded. **b,** Image of the DMS showing ChR2-eGFP expression in dMSNs (cell membrane) and eGFP expression in CINs (cytoplasm). **c**, Picrotoxin blocked oIPSCs (470 nm, 5 ms) recorded in DMS CINs (left); summarized oIPSC data (right); n = 8/3. **d,** A burst of light stimulation (20 Hz, 5 pulses) of SNr-projecting dMSNs inhibited CIN firing. Representative trace (top), repeated for multiple sweeps (middle), and the corresponding peristimulus histogram (bottom). **e**, Rabies-GFP was infused into the DMS of ChAT-Cre;D1tdT mice after helper virus infusion (TVA, RG). dMSNs presynaptic to the starter CINs, labeled with GFP and tdT, were yellow. **f-h**, Representative images of the GPi showing green (GFP^+^) fibers from putative presynaptic dMSNs (f), red fibers (tdT^+^) from dMSNs (g), and their overlap (h). **i-k**, Representative images of the SNr showing green fibers from putative presynaptic dMSNs (i), red fibers from dMSNs (j), and their overlap (k). Scale bars: 15 µm (b); 10 ms, 20 pA (c); 100 µm (f-h); 1 mm (i-k).

## DISCUSSION

The present study has identified a striatal mechanism underlying the reduction in cognitive flexibility induced by the reinforcement of substance use. We found that cocaine administration impaired cognitive flexibility, as evidenced by deficits in reversal learning of a two action-outcome instrumental task. Additionally, cocaine administration attenuated the activity of DMS CINs, a change that was associated with an increase in the inhibitory presynaptic inputs onto these neurons. Furthermore, we discovered that dMSNs provided the primary inhibitory inputs to DMS CINs. We found that cocaine administration potentiated glutamatergic inputs to DMS dMSNs and that cocaine or alcohol administration enhanced inhibitory dMSN→CIN connectivity. Moreover, chemo- or optogenetic inhibition of DMS CINs reduced reversal and extinction learning. Importantly, we discovered that SNr-projecting dMSNs, which mediate reinforcement, also sent axonal collaterals to inhibit DMS CINs, which mediate cognitive flexibility. These findings imply that the local inhibitory dMSN→CIN circuit mediates a reinforcement-induced reduction in cognitive flexibility.

### Reinforcement, cognitive flexibility, and CIN activity

We found that experimenter-delivered cocaine impaired reversal learning in a two-action-outcome instrumental task. Although cocaine and alcohol have previously been shown to suppress reversal learning (Jentsch *et al*., 2002; Obernier et al., 2002), cocaine-induced deficits in reversal learning have not previously been reported using an instrumental task. We observed that while cocaine administration did not alter the initial learning of the action-outcome contingency task, cocaine-administered mice exhibited reduced nose-poking responses during reversal learning, when the contingencies were reversed. A previous study found a similar reduction following contingency reversal after intra-DMS infusion of a cholinergic M2/M4 receptor agonist (Bradfield *et al*., 2013). This reduction in nose-poking is not likely to reflect reduced locomotion since cocaine withdrawal has been shown to increase, rather than decrease, this activity (Mañas-Padilla et al., 2021). The reduction in nose-poking is also unlikely to result from a cocaine-induced decrease in motivation because both the cocaine and saline groups showed equal increases in their responses during initial learning as the reinforcement schedule became progressively more challenging (FR1→RR20).

We also found that both experimenter-delivered (non-contingent i.p.) and voluntary administration (contingent IVSA) of cocaine led to a long-lasting reduction in the spontaneous firing frequency of DMS CINs. This reduction is unlikely to be caused by a direct effect on D2Rs because long-term cocaine exposure reduces dopamine release and/or the availability of inhibitory D2Rs; these dopamine or D2R changes would be more likely to increase CIN firing (Ashok et al., 2017). Therefore, the reduction in DMS CIN firing is more likely to reflect changes in synaptic transmission. Interestingly, a recent study also reported a downregulation of CIN activity in the ventral striatum after acute amphetamine exposure (Ztaou et al., 2021), which is consistent with our concept that addictive substances suppress striatal CIN activity. Surprisingly, acute cocaine exposure *in vivo* or *ex vivo* has been shown to increase striatal cholinergic activity (Mark et al., 1999; Witten et al., 2010). However, the long-term effects of cocaine administration on CIN firing have not been reported previously. The opposite regulation of CIN firing by acute and chronic cocaine exposure implies that cocaine withdrawal could lead to adaptations in CIN firing.

### Reinforcement and the MSN→CIN circuit in the DMS

Consistent with the reduction in CIN firing activity, we discovered that cocaine or alcohol administration enhanced GABAergic transmission in CINs. This enhancement results from elevated presynaptic GABA release, as indicated by a decreased paired-pulse ratio and an increased sIPSC frequency. Our analysis of the source of these GABAergic inputs using rabies-mediated retrograde tracing revealed that the majority of striatal neurons presynaptic to CINs were D1R-positive. In the striatum, GABAergic D1R-expressing dMSNs have been shown to synapse onto CINs (Chuhma et al., 2011; Lim and Surmeier, 2020). Additionally, striatal tyrosine hydroxylase interneurons also express D1Rs and send GABAergic inputs to DMS CINs (Dorst *et al*., 2020; English et al., 2011; Holley et al., 2015). However, there are far fewer interneurons than dMSNs. CINs are also known to receive GABAergic inputs from the external globus pallidus (Klug et al., 2018), but pallidus neurons do not express D1Rs (Lu et al., 2021b). Consequently, slice recordings confirmed that dMSNs send more robust GABAergic inputs to CINs than do iMSNs. This characterization of dMSN versus iMSN GABAergic inputs to CINs has not been reported previously. However, earlier studies noted that there were more substance P (co-released by dMSNs)-positive synapses than enkephalin (co-released by iMSNs)-positive synapses onto CINs (Bolam et al., 1986; Martone et al., 1992). These observations are consistent with our findings indicating that dMSNs provide the primary GABAergic inputs to DMS CINs.

GABAergic dMSN→CIN transmission was enhanced by the administraton of an addictive substance and by intracranial dMSN self-stimulation. Since we did not observe any postsynaptic changes in GABA_A_R responsiveness, the observed enhancement of dMSN→CIN transmission likely resulted from a potentiation of dMSN activity. Consistent with this, cocaine administration increased AMPA-induced currents and the AMPAR/NMDAR ratio in DMS dMSNs. This cocaine-induced potentiation of dMSN activity has previously been reported in the nucleus accumbens (Terrier et al., 2015) but not in the DMS. Interestingly, alcohol administration, intracranial self-stimulation of dopamine neurons, and self-administration of natural rewards have all been shown to potentiate striatal dMSN activity by increasing AMPAR trafficking (Cheng *et al*., 2017; Kravitz *et al*., 2012; Luscher, 2016; Pascoli *et al*., 2015; Shan *et al*., 2014; Wang et al., 2012; Wang et al., 2015). Taken together, these findings indicate that reinforcement of addictive substance use potentiates dMSN activity, thus strengthening GABAergic dMSN→CIN transmission, which reduces CIN activity.

### DMS CIN-mediated regulation of cognitive flexibility

We found that chemogenetic inhibition of DMS CINs compromised reversal learning. We also found that optogenetic inhibition of CIN activity, specifically on reward delivery, impaired reversal and extinction learning. However, inhibiting CIN activity during lever presses had no effect. Interestingly, magazine entries increased when the task schedule changed, and declined with continued training. This observation is consistent with a previous report that novice mice showed significantly more magazine entries than did well-trained mice (Matamales *et al*., 2020). Novel action-outcome associations are established by repeated checking the outcome and comparing it with the expected reward. When the action-outcome contingency changes (reversal learning), the animals increase their magazine entries to confirm this change. In support of our observations, a previous study showed that ACh activity in the nucleus accumbens transiently increased after reward delivery, but only on the first day of association learning; on day 5, this increase in ACh activity was absent (Al-Hasani et al., 2021). Similar observations of specific increases in ACh activity in the amygdala during salient events have also been reported. The absence of any difference in nose-poking between study groups upon contingency reversal imply that DMS CIN inhibition had no effect on motivation or locomotor activity, consistent with previous reports (Bradfield and Balleine, 2017; Bradfield *et al*., 2013). Therefore, our study provides evidence that DMS CIN activity during reward delivery mediates behavioral flexibility.

In summary, we demonstrate that cocaine-mediated reinforcement induces long-lasting facilitation of inhibitory dMSN→CIN transmission, thus reducing cognitive flexibility in a two action-outcome reversal learning task. This cocaine-mediated deficit of cognitive flexibility is associated with an increased GABAergic input, which reduces DMS CIN activity. Chemo- or optogenetic inhibition of DMS CINs, conducted to mimic the reinforcement effect of addictive substances, also reduced cognitive flexibility. Importantly, SNr-projecting dMSNs, which mediate reinforcement, sent axonal collaterals to inhibit DMS CINs, which mediate behavioral flexibility. Our findings demonstrate that the local inhibitory dMSN→CIN circuit mediates the reinforcement-induced reduction in cognitive flexibility.

## Materials and Methods

### Reagents

Cocaine solutions were prepared with (-)-cocaine hydrochloride (NIDA Drug Supply Program; cat. #9041-001) in saline. CNO was obtained from the NIMH chemical synthesis and drug supply program. DNQX disodium salt (AMPA/kainate antagonist) were obtained from Abcam (ab120169). AP5 was obtained from R&D systems. All other reagents was obtained from Sigma. For behavioral experiments, food pellets were obtained from Bioserve and sucrose was obtained from Tocris. Further details regarding viruses and animal breeding is provided in the Supplementary Information.

### Stereotaxic Virus Infusions

Animals were anesthetized with 3-4% (vol/vol) isoflurane at 1.0 L/min and placed on a heating pad to maintain their body temperature. The head was leveled, and craniotomies were performed using stereotaxic coordinates from the mouse (Franklin and Paxinos, 2007) or rat (Paxinos and Watson, 2007) brain atlas. Depending on the experiment, viruses were infused using injectors at the following coordinates from Bregma. Rat behavior experiments (Figure 5h-j, 7): DMS1-AP: 0.48, ML: ±2.5, DV: −4.7, DMS2-AP: −0.24, ML: ±3, DV: −4.7. Mouse behavior experiments (Figure 6) DMS1-AP: 0.38, ML: ±1.55, DV: −2.9, DMS2-AP: 0.02, ML: ±1.8, DV: −2.7. Mouse electrophysiology and histology experiments: DMS-AP: 0.38, ML: ±1.55, DV: −2.9, SNr-AP: −3.28, ML: ±1.2, DV: −4.5. For rat experiments, 0.8-1 µL of the virus was infused at each injection site at a rate of 0.1 µL/min. Injectors were kept in place for 10 min to allow the virus to diffuse. For mouse virus infusions, 0.4-0.5 µL of the virus was infused at each injection site. Animals were then returned to their home cages for 6 weeks before being used in experiments (Cheng *et al*., 2017; Ma *et al*., 2018). For Figure 3, rabies-GFP was infused into the DMS with the injectors at a 10-degree angle, to avoid contamination of the infusion track. For Figure 7, one cohort of rats began initial learning of the two action-outcome reversal learning task two weeks after viral infusions. When the rats reached the RR20 training sessions of initial learning, optical fibers were implanted. One week after fiber implantations, the rats resumed behavioral testing.

### Optical Fiber Implantations

Rats were anesthetized using 2-3% (vol/vol) isoflurane and were mounted on a stereotaxic frame (Kopf instruments). An incision was made, and bilateral optical fiber implants (300-nm core fiber secured to a 2.5-mm ceramic ferrule with 5 mm of fiber extending past the end of the ferrule) were lowered into the DMS at an angle of 5 degrees at the coordinates of AP: 0.12, ML: ±3.2, DV: −4.55 from Bregma. Implants were secured to the skull using metal screws and dental cement (Henry Schein), and covered with denture acrylic (Lang Dental). The incision was closed around the head cap and the skin was vet-bonded to the head cap. Rats were monitored for 1 week or until they had resumed normal activity. The methods were adapted from (Ma *et al*., 2018).

### Jugular Vein Catheter Implantation Surgery

The jugular vein catheter was constructed in-house, with the main parts purchased from SAI Infusion Technologies. Prior to the surgery, all catheters were tested for leaks using sterile water, and the intracardial tubing end was trimmed to 1.2 mm from the anchoring bead. Catheter implantation surgery was performed under oxygen/sevoflurane vapor and based on protocols described in Thomsen & Saine (Thomsen and Caine, 2007). Briefly, the catheter base was placed above the mid-scapular region, and the intracardial tubing was passed subcutaneously over the right shoulder and inserted 1.0-1.2 mm into the jugular vein. Two suture threads were tied around the vein above and below the anchoring beads to secure the intracardial tubing. Following surgery, mice were rested for 24 h. They received daily administration of heparin and cefazolin solution (30 USP units/mL heparin and 67 mg/mL cefazolin in 0.02 mL of 0.9% saline) through the catheter for 7 d. Antibiotic ointment was applied daily to the skin at the surgery site. Thereafter, catheters were flushed daily with 0.03 mL heparinized saline (0.9%) throughout the self-administration sessions. Catheter patency was examined weekly by using it for injection of 0.03 mL of 15 mg/mL ketamine and 0.75 mg/mL midazolam, with loss of righting reflex required within 5 sec (Huebschman et al., 2021).

### Slice Electrophysiology

Coronal sections of the dorsomedial striatum (thickness: 250 µm) were cut at a speed of 0.14 mm/s in ice-cold cutting solution containing (in mM) 40 NaCl, 148.5 sucrose, 4.5 KCl, 1.25 NaH_2_PO_4_, 25 NaHCO_3_, 0.5 CaCl_2_, 7 MgSO_4_, 10 dextrose, 1 sodium ascorbate, 3 myo inositol and 3 sodium pyruvate, and bubbled with 95% O_2_ and 5% CO_2_. The slices were then incubated for 45 min in a 1:1 mixture of cutting and external solution containing (in mM) 125 NaCl, 4.5 KCl, 2.5 CaCl_2_, 1.3 MgSO_4_, 1.25 NaH_2_PO_4_, 25 NaHCO_3_, 15 sucrose and 15 glucose, and saturated with 95% O_2_ and 5% CO_2_.

Potassium-based intracellular solution was used for all cell-attached and current-clamp recordings in Figure 1, 4h-k, 8k and Supplementary Figure 2; this contained (in mM) 123 potassium gluconate, 10 HEPES, 0.2 EGTA, 8 NaCl, 2 MgATP and 0.3 NaGTP, with the pH adjusted to 7.3 using KOH. Cesium intracellular solution was used for all other recordings; this contained (in mM) 119 CsMeSO_4_, 8 TEA.Cl, 15 HEPES, 0.6 ethylene glycol tetraacetic acid (EGTA), 0.3 Na_3_GTP, 4 MgATP, 5 QX-314.Cl and 7 phosphocreatine, with the pH adjusted to 7.3 using CsOH.

Cell-attached recordings were performed in a bath containing the external solution. NBQX (10 µm) and AP5 (50 µm) were used to block all glutamatergic transmission during IPSC recordings (Figure 2, 4, 5a, 5f-g, 5j, 8j, Supplementary 3). Picrotoxin (100 µm) was used to block all GABAergic transmission for EPSC recordings (Figure 5c-d). The recording bath was maintained at 32°C, and the perfusion speed was set to 2-3 mL/min. CINs in slices were identified by their labeled color, large size, and their spontaneous firing. The holding potential for CINs was −60 mV. D1 MSNs were held at −70 mV to measure AMPA currents and at +40 mV to measure NMDA currents (Fig 5F-I).

Spontaneous CIN firing was recorded for 5 min and the average frequency was then calculated. sIPSCs were recorded for 2 min for calculation of the average sIPSC frequency and amplitude. For eIPSC recordings, the electrical stimulation electrode was placed 100-150 µm away from the CIN of interest. In order to determine the paired-pulse ratios, two electrical stimulations separated by 200 ms were applied and the ratio of peak CIN eIPSC response 2 over peak response 1 was calculated. To record optically-induced IPSCs in CINs following dMSN stimulation, we delivered a 5-ms flash of light at 470 nm for ChR2 and at 590 nm for Chrimson. To record the effect of burst dMSN stimulation on CIN firing, optical stimulation at 470 nm (20 Hz, 5 pulses, 5 ms) was used for multiple sweeps. In a single sweep, the CIN was allowed to fire spontaneously for 10 sec, followed by train dMSN stimulation for 500 ms. Peristimulus histograms were generated by dividing the sweep time into bins of 1 sec (Fig. 4h) or 0.5 sec (Fig. 4k). To compare dMSN→CIN connectivity in animals exposed to alcohol, cocaine or intracranial dMSN self-stimulation versus controls, we followed a similar strategy; however, we recorded the oIPSCs at multiple stimulation intensities to generate an input-output (I-O) curve. To measure AMPA-induced currents in dMSNs, AMPA (5 µM) was bath-applied to the slices for 15 sec, and holding currents were recorded at a frequency of 0.2 Hz. To measure AMPA/NMDA ratios in dMSNs, the peak AMPAR current (recorded at a holding potential of −70 mV) was divided by the peak NMDAR current (recorded at a holding potential of +40 mV). Because A2A-Cre;Ai167;ChATeGFP mice were prone to early death, recordings were performed on postnatal days 27-30 in these mice. When recordings were made on the 24^th^ day after cocaine withdrawal, this time point was chosen to align with the first day of reversal testing (Fig. 1f). The 24 days included 3 days of food restriction, 4 magazine training sessions, 11 initial learning sessions, and 6 weekends.

### Behavior

Behavioral experiments were conducted in 4- to 7-month-old animals that were housed at 23°C with a 12-h light:dark cycle. All behavioral experiments were performed during the dark phase. Mice were housed singly, while rats were housed in pairs for behavioral experiments.

### Cocaine-induced Locomotor Sensitization

Saline was injected intraperitoneally (i.p.) in wild-type mice for 3 d to habituate them to the injection procedure and open field locomotion testing. After habituation, half of the animals continued to receive saline, while the other half received daily i.p. cocaine injections over the next 7 d. After each injection, the mice were placed in an open field chamber and locomotion was measured for 30 min. Any locomotor activity resulted in beam breaks, which were detected and recorded by the software (Kinder Scientific, CA).

### Reversal Learning of Instrumental Tasks in Mice (4 Sub-Sessions)

This procedure was adapted from (Bradfield and Balleine, 2017).

#### Apparatus design

The operant chambers (Coulbourn Instruments, MA) contained a house light, a white noise speaker, and two nose-poke holes (one on the left wall and the other on the right wall). Similarly, it had two magazines for reward delivery (a food magazine on the left wall and a sucrose magazine on the right wall). The training protocols were written in Graphic State, and the chambers were controlled using Graphic State RT.

#### Food deprivation

All mice were food-deprived for at least 3 d before the start of training to bring their respective bodyweights down to 80% of their free weight. Bodyweight was then maintained at 80% throughout the whole experiment.

#### Magazine training (4 d)

Mice were initially trained to enter the food or sucrose magazines to collect either a food pellet (20 mg; Dustless Precision Pellets Rodent, Grain-Based; Bio-Serv) or sucrose solution (20% solution in drinking water, 5-sec availability), respectively. No nose-pokes were available for poking during magazine training. Each session started with the illumination of the house light. Twenty deliveries of either food pellets or sucrose solution were made in the respective magazine using the RT60 schedule, where the average time between deliveries was 60 sec. Only one type of reward (either food or sucrose) was delivered during each session. The reward type was distributed evenly over the four magazine training sessions. A value of 200 total magazine entries was set as the threshold for sufficient motivation in these mice, before moving on to the instrumental training phase. Each mouse used the same operant chamber throughout all their training sessions.

#### Initial training (11 d)

Each training session had 4 sub-sessions (2 food and 2 sucrose sub-sessions in a random order) separated by a 2.5-min timeout period when the house light was turned off. During each sub-session, only one type of reward (food or sucrose) was available, and mice were required to nose-poke to receive reward delivery. A sub-session ended either when the mice were able to acquire 10 rewards or when 10 min of the sub-session had elapsed, whichever occurred first. The associations were: left nose-poke (A1) for food delivery (O1, left magazine) and right nose-poke (A2) for sucrose delivery (O2, right magazine). For the first 2 d, mice were trained on the FR1 schedule, where one nose-poke resulted in one reward delivery. Subsequently, the mice were trained on RR5 (p = 0.2 for 3 d), RR10 (p = 0.1 for 3 d) and RR20 (p = 0.05 for 3 d) schedules.

#### Reversal training (4 d)

This was conducted as described for the initial training (above), except that nose-poking on the left (A1) resulted in sucrose reward delivery (O2), and nose-poking on the right (A2) resulted in food reward delivery (O1). The training employed the RR20 (p = 0.2) schedule for 4 d.

### Cocaine Intravenous Self-administration

Operant chambers (Coulbourn Instruments, MA) containing two nose-poke ports were each housed in light- and sound-attenuating chambers. The chambers had cue lights inside and above each port and a cannula that ran through the ceiling of the chamber could be attached to the implanted jugular vein catheter. Mice were trained to self-administer an intravenous infusion of cocaine (1 mg/kg/infusion in sterile 0.9% saline). The drug infusion volume was 0.56 mL/kg and the concentration was 0.18 mg/mL. The length of each infusion was calculated using weight (kg)/0.01 sec. The experiment ran for 7 d, with one 3-h session per day. The maximum number of cocaine infusions that the animals could receive during each session was 30. Illumination of the house light signaled reward availability and the light was turned off following each infusion. There was a 20-sec timeout period following each infusion, during which an effective nose-poke did not trigger an infusion. The acquisition phase was conducted with an FR1 schedule, where each effective nose-poke on the active port resulted in one drug infusion coupled with activation of the cue light inside the nose-poke hole (Huebschman *et al*., 2021).

### Intracranial Optogenetic Self-Stimulation of dMSNs

Rats were connected to the laser via patch cables before they were placed in an operant chamber containing two nose-poke holes with cue lights above them. An active nose-poke resulted in activation of the corresponding cue light (10 pulses; 0.5-sec ON and 0.5-sec OFF); this was paired with laser stimulation (30 trains of burst stimulation at an inter-burst interval of 250 ms) that occurred after a 5-sec delay (Pascoli *et al*., 2015). Each burst included 5 pulses (470 nm, 4 ms) at 20 Hz. An inactive nose-poke did not trigger the cue light or laser stimulation. Nose-poking during laser stimulation had no additional effect, but these nose-pokes were recorded. Each session lasted for 30 min. Rats were initially trained on an FR1 schedule. Rats that showed a preference for the active nose-poke were then moved to the FR2 schedule for 3 d, followed by the FR3 schedule for 4 d. All experiments were performed in the dark cycle and rats were food-deprived to increase their performance and activity.

### Reversal Learning of an Instrumental Task in Mice (2 Sub-Sessions)

The food deprivation, chamber design, and magazine training procedures used were similar to those described in the “Reversal Learning of Instrumental Tasks in Mice (4 Sub-Sessions)” section (above). The methods were adapted from (Bradfield *et al*., 2013; Matamales *et al*., 2020).

#### Initial learning (11 d)

Each training session had two sub-sessions (1 food and 1 sucrose sub-session in random order), separated by a 5-min timeout period. During each sub-session, only one type of reward (food or sucrose) was available; mice were required to nose-poke to receive the reward delivery, while the other nose-poke area was covered. A sub-session ended either when the mice were able to acquire 20 rewards or when 30 min had elapsed, whichever occurred first. After a 5-min timeout period, the animals were trained for the other reward. The associations used were: left nose-poke (A1) for food delivery (O1, left magazine) and right nose-poke (A2) for sucrose delivery (O2, right magazine). Mice were trained on an FR1 schedule for the first 2 d and then moved to RR5 (p = 0.2 for 3 d), RR10 (p = 0.1 for 3 d) and RR20 (p = 0.05 for 3 d) schedules. This was followed by two sessions of devaluation testing.

#### Devaluation testing (2 d)

To induce satiety, mice were given free access to either food or sucrose (selected randomly) for 1 h prior to testing. A separate new cage was used for this and it was placed in a different room to where the instrumental training had been conducted. After feeding, the mice were placed in the operant chamber. The house light was turned on and their total numbers of left or right nose-pokes were recorded for 10 min, with neither resulting in reward delivery. This was repeated on the following day, except that the mice were pre-fed with the other reward.

#### Reversal learning (4 d)

This was conducted as described for the initial training (see above), except that nose-poking on the left (A1) resulted in sucrose reward delivery (O2), and nose-poking on the right (A2) resulted in food reward delivery (O1). The training employed the RR20 (p = 0.2) schedule for 4 d and was followed by devaluation testing, as described above.

### Reversal Learning of an Instrumental Task in Rats (4 Sub-Sessions)

This procedure was adapted from (Bradfield and Balleine, 2017) and only differed from the methods specified in the “Reversal Learning of Instrumental Tasks in Mice (4 Sub-Sessions)” in three respects: a) rats were used instead of mice; b) rat operant chambers were used; and c) instrumental behavior was measured using lever presses rather than nose-pokes. The procedures for magazine training, initial training, reversal training, and devaluation testing remained the same.

#### Apparatus

Each operant chamber had two separate magazines located in the center of the right wall; these were connected to a sucrose dipper (left) or a food dispenser (right). On either side of the magazines, there was a retractable lever (left and right). The sensors for both the food and sucrose magazine entries were the same, which meant that we were unable to differentiate between food and sucrose magazine entries.

### Extinction Learning

Rats were retrained on the operant instrumental task using the RR20 schedule (Fig. 7c) until they showed a stable level of performance for at least 2 d. For extinction training (3 sessions), we removed the rewards from the operant setup and kept all the other parameters the same. Optical stimulation (590 nm, 5 s) was provided to inhibit DMS CINs when the reward was supposed to be delivered.

### Intermittent-Access to 20% Alcohol Two-Bottle Choice Drinking Procedure

One bottle containing drinking water and one bottle containing 20% alcohol were introduced to the mouse home cages at noon on Mondays, Wednesdays, and Fridays. The alcohol bottle was replaced with a bottle containing only drinking water at noon on Tuesdays, Thursdays, and Saturdays. This schedule was maintained for 8 weeks, following which we measured the amount of alcohol each mouse drank per day. Electrophysiology recordings from alcohol-exposed mice were performed 1 d after the last alcohol exposure. The methods were adopted from (Cheng *et al*., 2017).

### Histology

Mice and rats were anesthetized and perfused intracardially with 4% paraformaldehyde (PFA) in phosphate-buffered saline (PBS). The brains were then extracted and submerged in 4% PFA/PBS solution for one day at 4°C, following which they were transferred to a 30% sucrose solution in PBS. Once the brains completely sank in the sucrose solution, they were cut into 50-µm thick sections using a cryostat. The slices were stored in a PBS bath at 4°C prior to mounting on slides for imaging using a confocal laser-scanning microscope (Fluoroview-1200, Olympus). All images were processed using Imaris 8.3.1 (Bitplane, Zurich, Switzerland). ChAT-positive neurons were identified by staining with the anti-ChAT antibody (AB144P) and labeling with far-red fluorescence (647 nm) using a donkey anti-goat antibody (A21447).

### Data Analysis and Statistics

Each experiment was repeated in 3-7 animals and the resulting data were pooled for statistical analysis. The minimum sample size requirement for electrophysiology recordings was set at 3 (animals) and 8 (neurons). Outliers and unstable recordings with a change in series resistance of more than 10% were excluded from the analyses. For behavioral experiments, the control and experimental groups were age-matched. Data collection was randomized. No data were excluded, unless stated otherwise. All data are expressed as mean ± s.e.m. Statistical analyses were performed using Sigma Plot software. Normal distribution was tested and unpaired *t* test, paired *t* test, or two-way RM ANOVA followed by *post-hoc* Tukey test used to determine statistical significance as appropriate, with an alpha value of 0.05. All animal care and experimental procedures were approved by the Texas A&M University Animal Care and Use Committee.

## Acknowledgements

This research was supported by NIAAA R01AA027768 (JW), U01AA025932 (JW) and NIDA R01DA046457 (RJS).

## Author contributions

J.W. and H.G. conceived the project and designed the experiments. H.G., X.X., Y.C. and Y.H. performed the behavioral experiments and analyzed the corresponding data. H.G., J.L. and X.Z. performed the electrophysiological experiments and analyzed the corresponding data. Y.C., X.W., H.G. and J.W. performed the histology experiments. X.W. performed animal breeding for all experiments. J.W., H.G. and A.E. wrote the manuscript with substantial inputs from R.J.S. and L.N.S.

## Supplementary Information

### Supplementary Figures

**Supplementary Figure 1.**
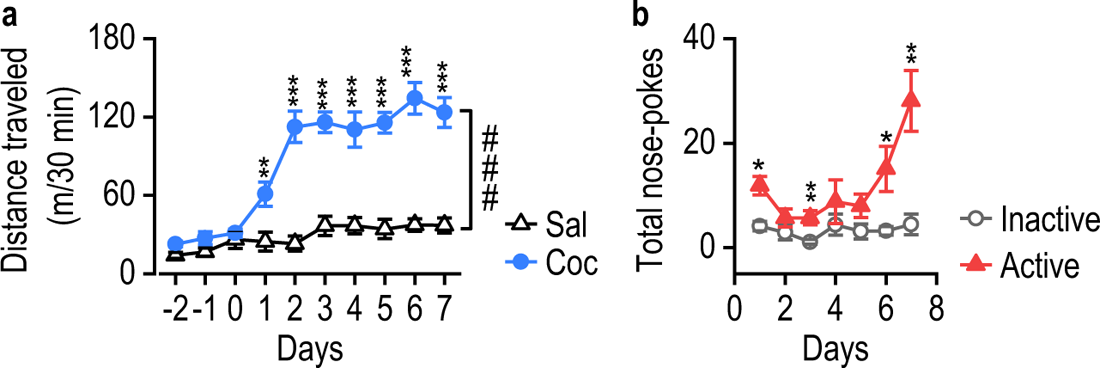
**a**, Repeated cocaine injections caused locomotor sensitization in ChAT-eGFP mice; ^###^*p* < 0.001 for the indicated comparison, two-way RM ANOVA; *\*\*\*p* < 0.001 and \*\**p* < 0.01 versus the saline group on the same days, Tukey *post-hoc* test, n = 5 mice (Sal) and 5 mice (Coc). **b**, Total active versus inactive nose-pokes to receive intravenous cocaine infusions (1.5 mg/kg/infusion) across training sessions in ChAT-eGFP mice; \*\**p* < 0.01, \**p* < 0.05 for the indicated training sessions, unpaired *t* test, n = 8 mice.

**Supplementary Figure 2.**
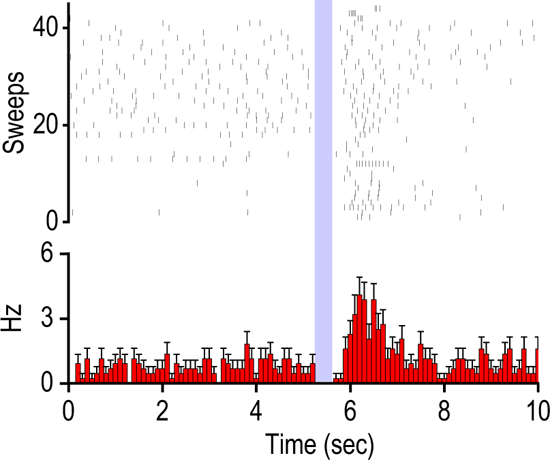
Burst stimulation (20 Hz, 5 ms, 5 pulses) of DMS dMSNs caused pause-rebound CIN firing in D1-Cre;Ai32 mice. Top, time course of 44 sweeps from 8 neurons; bottom, corresponding peristimulus histogram.

**Supplementary Figure 3.**
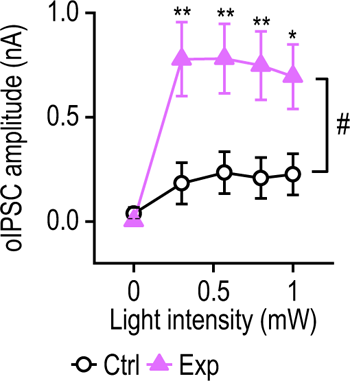
Summarized data comparing oIPSC^dMSN→CIN^ amplitudes at multiple stimulation intensities in DMS slices from rats that underwent optogenetic intracranial self-stimulation of dMSNs (Exp) and from control rats that did not self-stimulate dMSNs (Ctrl); ^#^*p* < 0.05 for the indicated comparison, two-way RM ANOVA; \*\**p* < 0.01, \**p* < 0.05 versus the Ctrl group at the same stimulating intensity, *post-hot* Tukey test; n = 5, 5 (Ctrl) and 4, 1 (Exp).

**Supplementary Figure 4.**
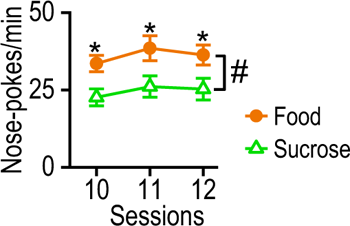
**a**, There was greater motivation to nose-poke for food than for sucrose, with significantly more nose-pokes per min for food than for sucrose during RR20 sessions in all mice; ^#^*p <* 0.05 by two-way RM ANOVA followed by Tukey *post hoc* test; \**p* < 0.05 versus the sucrose group on the same session by Tukey *post hoc* test, n = 16 mice.

**Supplementary Figure 5.**
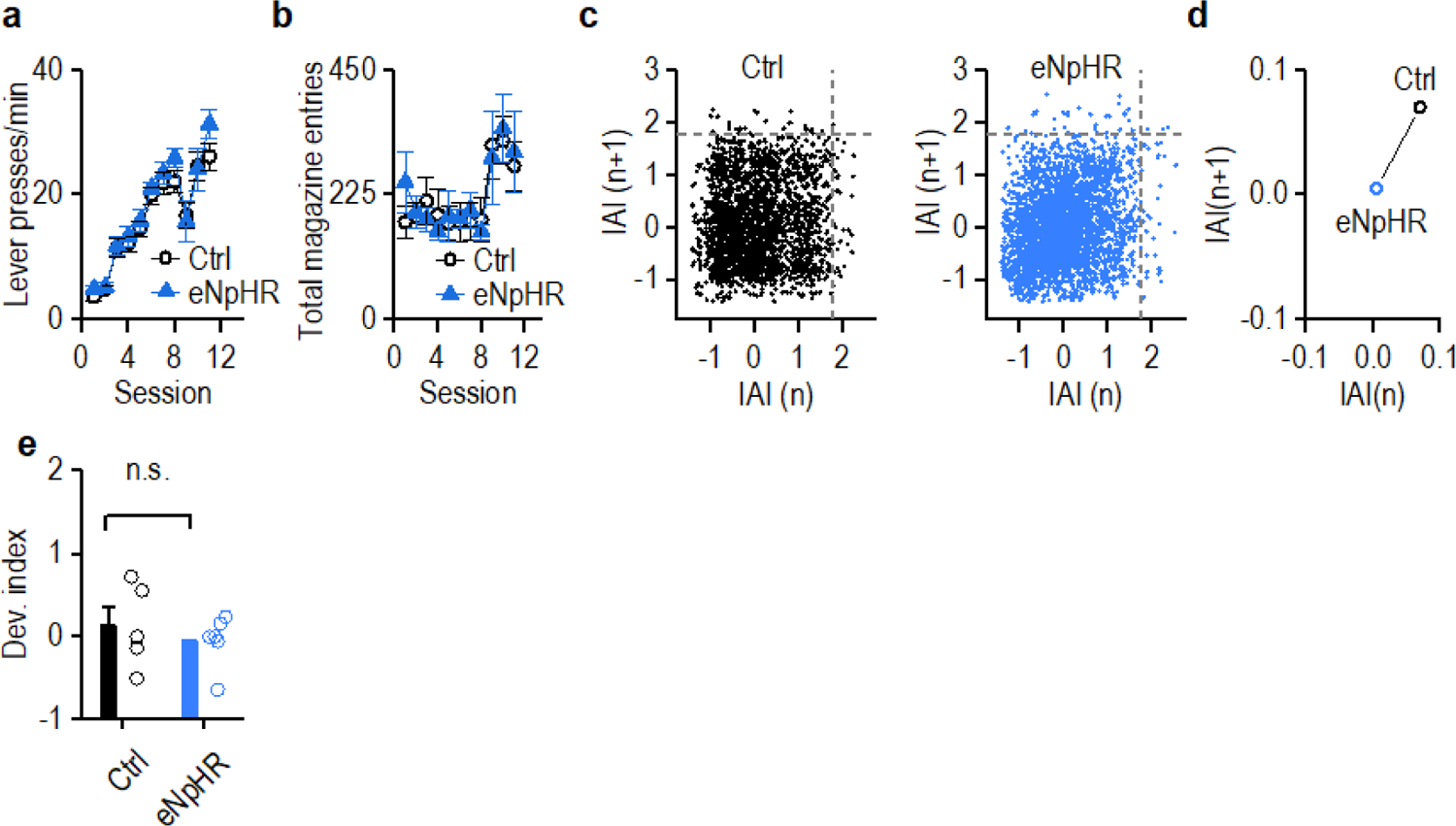
**a**, Lever presses across sessions for the control and experimental groups during initial training; n = 9 (Ctrl) and 8 (Exp) rats. **b**, Total magazine entries across sessions for the control and experimental groups during initial training; n = 9 (Ctrl) and 8 (Exp) rats. **c**, Return maps of inter-action intervals (IAI) between magazine entries for the control and experimental groups on the first day of reversal learning (pooled group data). Each data point (x,y) represents the time difference between the previous (x) and the next time point of magazine entry (y). **d**, Plot illustrating the centers of the data points in (c) for the control and experimental groups. **e,** No difference in devaluation index for the 2nd devaluation was observed between the control and experimental groups when CIN inhibition was paired to lever presses, rather than reward deliveries, during reversal learning; n.s., not significant; n = 5 (Ctrl) and 6 (Exp) rats.

**Supplementary Figure 6.**
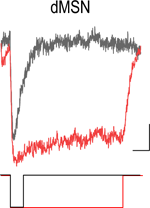
Functional verification of ChR2 expression on SNr-projecting dMSNs in the DMS. Longer optical stimulation (470 nm, 200 ms) generated a longer response than a shorter stimulation (470 nm, 5 ms), confirming direct depolarization. Scale: 30 ms, 30 pA.

### Supplementary Tables

**Table 1:**
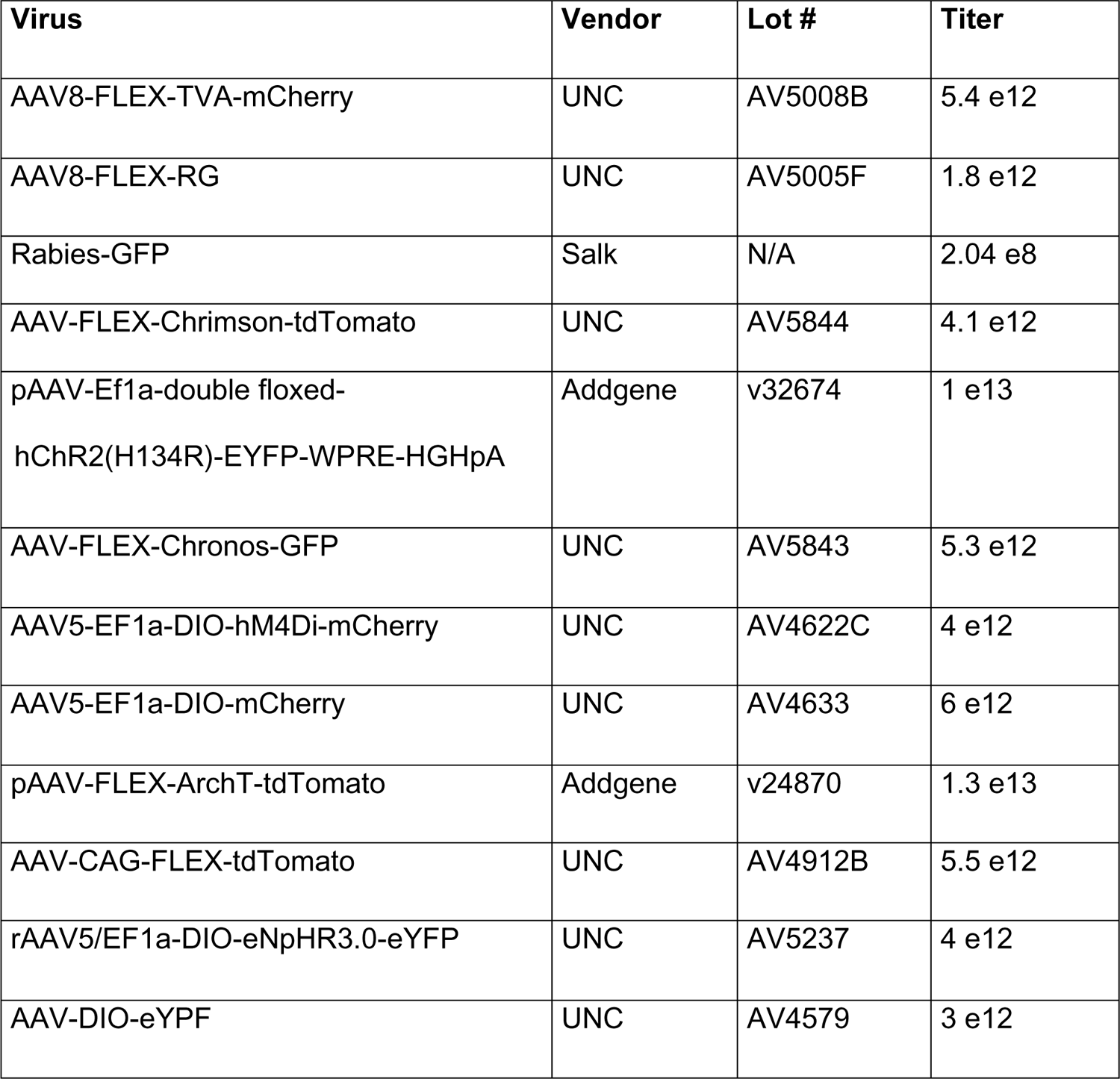
Virus Information

**Table 2:** Mouse Information

| ID | Genotype | Vendor | Lot # |
| --- | --- | --- | --- |
| 1 | C57BL/6J | Jackson Laboratories | 00664 |
| 2 | ChAT-Cre(+/+) | Jackson Laboratories | 028861 |
| 3 | deltaneoChAT-Cre(+/+) | Jackson Laboratories | 031661 |
| 4 | ChATeGFP(+/+) | Jackson Laboratories | 007902 |
| 5 | Ai14(+/+) | Jackson Laboratories | 007914 |
| 6 | D1tdTomato(+/-) | Jackson Laboratories | 016204 |
| 7 | Ai32(+/+) | Jackson Laboratories | 012569 |
| 8 | Drd1-Cre(+/-) | MMRRC | 29178 |
| 9 | Ai167(+/-) | Allen Brain Institution | N/A |
| 10 | A2A-Cre(+/-) | MMRRC | 036158-UCD |

**Table 3:** Rat Information

| ID | Genotype | Vendor | Lot # |
| --- | --- | --- | --- |
| 1 | D1-Cre(+/-) | RRRC | 856 |
| 2 | ChAT-Cre(+/-) | RRRC | 658 |
| 3 | tdTomato/Hom-KI(+/+) | Envigo Horizon | TGRL9660 |

## References

1. Al-Hasani, R., Gowrishankar, R., Schmitz, G.P., Pedersen, C.E., Marcus, D.J., Shirley, S.E., Hobbs, T.E., Elerding, A.J., Renaud, S.J., Jing, M., et al. (2021). Ventral tegmental area GABAergic inhibition of cholinergic interneurons in the ventral nucleus accumbens shell promotes reward reinforcement. Nature neuroscience, [Online ahead of print]. 10.1038/s41593-021-00898-2.

2. Aoki, S., Liu, A.W., Zucca, A., Zucca, S., and Wickens, J.R. (2015). Role of striatal cholinergic interneurons in set-shifting in the rat. J Neurosci 35, 9424–9431. 10.1523/JNEUROSCI.0490-15.2015.

3. Ashok, A.H., Mizuno, Y., Volkow, N.D., and Howes, O.D. (2017). Association of Stimulant Use With Dopaminergic Alterations in Users of Cocaine, Amphetamine, or Methamphetamine: A Systematic Review and Meta-analysis. JAMA Psychiatry 74, 511–519. 10.1001/jamapsychiatry.2017.0135.

4. Barker, J.M., Bryant, K.G., Osborne, J.I., and Chandler, L.J. (2017). Age and Sex Interact to Mediate the Effects of Intermittent, High-Dose Ethanol Exposure on Behavioral Flexibility. Front Pharmacol 8, 450. 10.3389/fphar.2017.00450.

5. Bolam, J.P., Ingham, C.A., Izzo, P.N., Levey, A.I., Rye, D.B., Smith, A.D., and Wainer, B.H. (1986). Substance P-containing terminals in synaptic contact with cholinergic neurons in the neostriatum and basal forebrain: a double immunocytochemical study in the rat. Brain Res 397, 279–289.

6. Bradfield, L.A., and Balleine, B.W. (2017). Thalamic control of dorsomedial striatum regulates internal state to guide goal-directed action selection. J Neurosci 37, 3721–3733. 10.1523/JNEUROSCI.3860-16.2017.

7. Bradfield, L.A., Bertran-Gonzalez, J., Chieng, B., and Balleine, B.W. (2013). The thalamostriatal pathway and cholinergic control of goal-directed action: interlacing new with existing learning in the striatum. Neuron 79, 153–166. 10.1016/j.neuron.2013.04.039.

8. Chen, B.T., Bowers, M.S., Martin, M., Hopf, F.W., Guillory, A.M., Carelli, R.M., Chou, J.K., and Bonci, A. (2008). Cocaine but not natural reward self-administration nor passive cocaine infusion produces persistent LTP in the VTA. Neuron 59, 288–297. 10.1016/j.neuron.2008.05.024.

9. Cheng, Y., Huang, C.C.Y., Ma, T., Wei, X., Wang, X., Lu, J., and Wang, J. (2017). Distinct synaptic strengthening of the striatal direct and indirect pathways drives alcohol consumption. Biological psychiatry 81, 918–929. 10.1016/j.biopsych.2016.05.016.

10. Chuhma, N., Tanaka, K.F., Hen, R., and Rayport, S. (2011). Functional connectome of the striatal medium spiny neuron. J Neurosci 31, 1183–1192. 31/4/1183 [pii]10.1523/JNEUROSCI.3833-10.2011.

11. Dong, Y., Taylor, J.R., Wolf, M.E., and Shaham, Y. (2017). Circuit and Synaptic Plasticity Mechanisms of Drug Relapse. J Neurosci 37, 10867–10876. 10.1523/JNEUROSCI.1821-17.2017.

12. Dorst, M.C., Tokarska, A., Zhou, M., Lee, K., Stagkourakis, S., Broberger, C., Masmanidis, S., and Silberberg, G. (2020). Polysynaptic inhibition between striatal cholinergic interneurons shapes their network activity patterns in a dopamine-dependent manner. Nat. Commun. 11, 5113. 10.1038/s41467-020-18882-y.

13. English, D.F., Ibanez-Sandoval, O., Stark, E., Tecuapetla, F., Buzsaki, G., Deisseroth, K., Tepper, J.M., and Koos, T. (2011). GABAergic circuits mediate the reinforcement-related signals of striatal cholinergic interneurons. Nat Neurosci 15, 123–130. 10.1038/nn.2984nn.2984 [pii].

14. Everitt, B.J., and Robbins, T.W. (2005). Neural systems of reinforcement for drug addiction: from actions to habits to compulsion. Nat Neurosci 8, 1481–1489.

15. Franklin, K.B.J., and Paxinos, G. (2007). The mouse brain in stereotaxic coordinates, 3 Edition (Academic Press).

16. Freeze, B.S., Kravitz, A.V., Hammack, N., Berke, J.D., and Kreitzer, A.C. (2013). Control of basal ganglia output by direct and indirect pathway projection neurons. J Neurosci 33, 18531–18539. 10.1523/JNEUROSCI.1278-13.2013.

17. Galaj, E., Kipp, B.T., Floresco, S.B., and Savage, L.M. (2019). Persistent Alterations of Accumbal Cholinergic Interneurons and Cognitive Dysfunction after Adolescent Intermittent Ethanol Exposure. Neuroscience 404, 153–164. 10.1016/j.neuroscience.2019.01.062.

18. Gerfen, C.R., and Surmeier, D.J. (2011). Modulation of striatal projection systems by dopamine. Annu Rev Neurosci 34, 441–466. 10.1146/annurev-neuro-061010-113641.

19. Gonzales, K.K., Pare, J.F., Wichmann, T., and Smith, Y. (2013). GABAergic inputs from direct and indirect striatal projection neurons onto cholinergic interneurons in the primate putamen. J Comp Neurol 521, 2502–2522. 10.1002/cne.23295.

20. Guo, Q., Wang, D., He, X., Feng, Q., Lin, R., Xu, F., Fu, L., and Luo, M. (2015). Whole-brain mapping of inputs to projection neurons and cholinergic interneurons in the dorsal striatum. PLoS One 10, e0123381. 10.1371/journal.pone.0123381.

21. Hellard, E.R., Binette, A., Zhuang, X., Lu, J., Ma, T., Jones, B., Williams, E., Jayavelu, S., and Wang, J. (2019). Optogenetic control of alcohol-seeking behavior via the dorsomedial striatal circuit. Neuropharmacology 155, 89–97. 10.1016/j.neuropharm.2019.05.022.

22. Holley, S.M., Joshi, P.R., Parievsky, A., Galvan, L., Chen, J.Y., Fisher, Y.E., Huynh, M.N., Cepeda, C., and Levine, M.S. (2015). Enhanced GABAergic inputs contribute to functional alterations of cholinergic interneurons in the r6/2 mouse model of Huntington’s disease. eNeuro 2, 0008–0014. 10.1523/eneuro.0008-14.2015.

23. Huebschman, J.L., Davis, M.C., Tovar Pensa, C., Guo, Y., and Smith, L.N. (2021). The fragile X mental retardation protein promotes adjustments in cocaine self-administration that preserve reinforcement level. Eur J Neurosci 54, 4920–4933. 10.1111/ejn.15356.

24. Hwa, L.S., Chu, A., Levinson, S.A., Kayyali, T.M., DeBold, J.F., and Miczek, K.A. (2011). Persistent escalation of alcohol drinking in C57BL/6J mice with intermittent access to 20% ethanol. Alcohol Clin Exp Res 35, 1938–1947. 10.1111/j.1530-0277.2011.01545.x.

25. Izquierdo, A., and Jentsch, J.D. (2012). Reversal learning as a measure of impulsive and compulsive behavior in addictions. Psychopharmacology (Berl) 219, 607–620. 10.1007/s00213-011-2579-7.

26. Jentsch, J.D., Olausson, P., De La Garza, R., 2nd, and Taylor, J.R. (2002). Impairments of reversal learning and response perseveration after repeated, intermittent cocaine administrations to monkeys. Neuropsychopharmacology 26, 183-190. 10.1016/S0893-133X(01)00355-4.

27. Kauer, J.A., and Malenka, R.C. (2007). Synaptic plasticity and addiction. Nat Rev Neurosci 8, 844–858.

28. Klug, J.R., Engelhardt, M.D., Cadman, C.N., Li, H., Smith, J.B., Ayala, S., Williams, E.W., Hoffman, H., and Jin, X. (2018). Differential inputs to striatal cholinergic and parvalbumin interneurons imply functional distinctions. Elife 7, e35657. 10.7554/eLife.35657.

29. Kravitz, A.V., Tye, L.D., and Kreitzer, A.C. (2012). Distinct roles for direct and indirect pathway striatal neurons in reinforcement. Nat Neurosci 15, 816–818. 10.1038/nn.3100nn.3100 [pii].

30. Kreitzer, A.C., and Malenka, R.C. (2008). Striatal plasticity and basal ganglia circuit function. Neuron 60, 543–554.

31. Lalive, A.L., Lien, A.D., Roseberry, T.K., Donahue, C.H., and Kreitzer, A.C. (2018). Motor thalamus supports striatum-driven reinforcement. Elife 7, e34032. 10.7554/eLife.34032.

32. Lim, S.A.O., and Surmeier, D.J. (2020). Enhanced GABAergic Inhibition of Cholinergic Interneurons in the zQ175(+/-) Mouse Model of Huntington’s Disease. Front Syst Neurosci 14, 626412. 10.3389/fnsys.2020.626412.

33. Lu, J., Cheng, Y., Xie, X., Woodson, K., Bonifacio, J., Disney, E., Barbee, B., Wang, X., Zaidi, M., and Wang, J. (2021a). Whole-brain mapping of direct inputs to dopamine D1 and D2 receptor-expressing medium spiny neurons in the posterior dorsomedial striatum. eNeuro 8, 0348–0320.2020. 10.1523/ENEURO.0348-20.2020.

34. Lu, J., Cheng, Y., Xie, X., Woodson, K., Bonifacio, J., Disney, E., Barbee, B., Wang, X., Zaidi, M., and Wang, J. (2021b). Whole-Brain Mapping of Direct Inputs to Dopamine D1 and D2 Receptor-Expressing Medium Spiny Neurons in the Posterior Dorsomedial Striatum. eNeuro 8. 10.1523/ENEURO.0348-20.2020.

35. Luscher, C. (2016). The emergence of a circuit model for addiction. Annu Rev Neurosci, 39:257–276. 10.1146/annurev-neuro-070815-013920.

36. Luscher, C., and Janak, P.H. (2021). Consolidating the circuit model for addiction. Annu Rev Neurosci, 44:173–195. 10.1146/annurev-neuro-092920-123905.

37. Luscher, C., and Malenka, R.C. (2011). Drug-evoked synaptic plasticity in addiction: from molecular changes to circuit remodeling. Neuron 69, 650–663. S0896-6273(11)00065-1 [pii]10.1016/j.neuron.2011.01.017.

38. Luscher, C., Robbins, T.W., and Everitt, B.J. (2020). The transition to compulsion in addiction. Nat Rev Neurosci 21, 247–263. 10.1038/s41583-020-0289-z.

39. Ma, T., Cheng, Y., Roltsch Hellard, E., Wang, X., Lu, J., Gao, X., Huang, C.C.Y., Wei, X., Ji, J., and Wang, J. (2018). Bidirectional and long-lasting control of alcohol-seeking behavior by corticostriatal LTP and LTD. Nat Neurosci 21, 373–383. 10.1038/s41593-018-0081-9.

40. Ma, T., Huang, Z., Xie, X., Cheng, Y., Zhuang, X., Childs, M., Gangal, H., Wang, X., Smith, L.N., Smith, R.J., et al. (2021). Chronic alcohol drinking persistently suppresses thalamostriatal excitation of cholinergic neurons to impair cognitive flexibility J Clin Invest Accepted.

41. Mañas-Padilla, M.C., Ávila-Gámiz, F., Gil-Rodríguez, S., Ladrón de Guevara-Miranda, D., Rodríguez de Fonseca, F., Santín, L.J., and Castilla-Ortega, E. (2021). Persistent changes in exploration and hyperactivity coexist with cognitive impairment in mice withdrawn from chronic cocaine. Physiol Behav 240, 113542. 10.1016/j.physbeh.2021.113542.

42. Mark, G.P., Hajnal, A., Kinney, A.E., and Keys, A.S. (1999). Self-administration of cocaine increases the release of acetylcholine to a greater extent than response-independent cocaine in the nucleus accumbens of rats. Psychopharmacology (Berl) 143, 47–53. 10.1007/s002130050918.

43. Martone, M.E., Armstrong, D.M., Young, S.J., and Groves, P.M. (1992). Ultrastructural examination of enkephalin and substance P input to cholinergic neurons within the rat neostriatum. Brain Res 594, 253–262.

44. Matamales, M., McGovern, A.E., Mi, J.D., Mazzone, S.B., Balleine, B.W., and Bertran-Gonzalez, J. (2020). Local D2- to D1-neuron transmodulation updates goal-directed learning in the striatum. Science 367, 549–555. 10.1126/science.aaz5751.

45. Matamales, M., Skrbis, Z., Hatch, R.J., Balleine, B.W., Gotz, J., and Bertran-Gonzalez, J. (2016). Aging-related dysfunction of striatal cholinergic interneurons produces conflict in action selection. Neuron 90, 362–373. 10.1016/j.neuron.2016.03.006.

46. McCracken, C.B., and Grace, A.A. (2013). Persistent cocaine-induced reversal learning deficits are associated with altered limbic cortico-striatal local field potential synchronization. J Neurosci 33, 17469–17482. 10.1523/JNEUROSCI.1440-13.2013.

47. Morris, G., Arkadir, D., Nevet, A., Vaadia, E., and Bergman, H. (2004). Coincident but distinct messages of midbrain dopamine and striatal tonically active neurons. Neuron 43, 133–143. 10.1016/j.neuron.2004.06.012.

48. Obernier, J.A., White, A.M., Swartzwelder, H.S., and Crews, F.T. (2002). Cognitive deficits and CNS damage after a 4-day binge ethanol exposure in rats. Pharmacol Biochem Behav 72, 521–532.

49. Pascoli, V., Hiver, A., Van Zessen, R., Loureiro, M., Achargui, R., Harada, M., Flakowski, J., and Luscher, C. (2018). Stochastic synaptic plasticity underlying compulsion in a model of addiction. Nature 564, 366–371. 10.1038/s41586-018-0789-4.

50. Pascoli, V., Terrier, J., Espallergues, J., Valjent, E., O’Connor, E.C., and Luscher, C. (2014). Contrasting forms of cocaine-evoked plasticity control components of relapse. Nature 509, 459–464. 10.1038/nature13257.

51. Pascoli, V., Terrier, J., Hiver, A., and Luscher, C. (2015). Sufficiency of Mesolimbic Dopamine Neuron Stimulation for the Progression to Addiction. Neuron 88, 1054–1066. 10.1016/j.neuron.2015.10.017.

52. Pascoli, V., Turiault, M., and Luscher, C. (2012). Reversal of cocaine-evoked synaptic potentiation resets drug-induced adaptive behaviour. Nature 481, 71–75. 10.1038/nature10709nature10709 [pii].

53. Paxinos, G., and Watson, C. (2007). The Rat Brain in stereotaxic coordinates, 6 Edition (Academic Press).

54. Prado, V.F., Janickova, H., Al-Onaizi, M.A., and Prado, M.A. (2017). Cholinergic circuits in cognitive flexibility. Neuroscience 345, 130–141. 10.1016/j.neuroscience.2016.09.013.

55. Ragozzino, M.E. (2007). The contribution of the medial prefrontal cortex, orbitofrontal cortex, and dorsomedial striatum to behavioral flexibility. Ann N Y Acad Sci 1121, 355–375. 10.1196/annals.1401.013.

56. Ragozzino, M.E., and Choi, D. (2004). Dynamic changes in acetylcholine output in the medial striatum during place reversal learning. Learn Mem 11, 70–77. 10.1101/lm.65404.

57. Roberts-Wolfe, D., Bobadilla, A.C., Heinsbroek, J.A., Neuhofer, D., and Kalivas, P.W. (2018). Drug Refraining and Seeking Potentiate Synapses on Distinct Populations of Accumbens Medium Spiny Neurons. J Neurosci 38, 7100–7107. 10.1523/JNEUROSCI.0791-18.2018.

58. Schultz, W. (1998). Predictive reward signal of dopamine neurons. J Neurophysiol 80, 1–27.

59. Shan, Q., Ge, M., Christie, M.J., and Balleine, B.W. (2014). The acquisition of goal-directed actions generates opposing plasticity in direct and indirect pathways in dorsomedial striatum. J Neurosci 34, 9196–9201. 10.1523/JNEUROSCI.0313-14.2014.

60. Simms, J.A., Steensland, P., Medina, B., Abernathy, K.E., Chandler, L.J., Wise, R., and Bartlett, S.E. (2008). Intermittent access to 20% ethanol induces high ethanol consumption in Long-Evans and Wistar rats. Alcohol Clin Exp Res 32, 1816–1823.

61. Sullivan, M.A., Chen, H., and Morikawa, H. (2008). Recurrent inhibitory network among striatal cholinergic interneurons. J Neurosci 28, 8682–8690. 10.1523/JNEUROSCI.2411-08.2008.

62. Terrier, J., Luscher, C., and Pascoli, V. (2015). Cell-Type Specific Insertion of GluA2-Lacking AMPARs with Cocaine Exposure Leading to Sensitization, Cue-Induced Seeking and Incubation of Craving. Neuropsychopharmacology 41, 1779–1789. 10.1038/npp.2015.345.

63. Thomsen, M., and Caine, S.B. (2007). Intravenous drug self-administration in mice: practical considerations. Behav Genet 37, 101–118. 10.1007/s10519-006-9097-0.

64. Ungless, M.A., Whistler, J.L., Malenka, R.C., and Bonci, A. (2001). Single cocaine exposure in vivo induces long-term potentiation in dopamine neurons. Nature 411, 583–587. 10.1038/35079077.

65. Wang, J., Ben Hamida, S., Darcq, E., Zhu, W., Gibb, S.L., Lanfranco, M.F., Carnicella, S., and Ron, D. (2012). Ethanol-mediated facilitation of AMPA receptor function in the dorsomedial striatum: implications for alcohol drinking behavior. J Neurosci 32, 15124–15132. 10.1523/JNEUROSCI.2783-12.2012.

66. Wang, J., Cheng, Y., Wang, X., Roltsch Hellard, E., Ma, T., Gil, H., Ben Hamida, S., and Ron, D. (2015). Alcohol elicits functional and structural plasticity selectively in dopamine D1 receptor-expressing neurons of the dorsomedial striatum. J Neurosci 35, 11634–11643. 10.1523/JNEUROSCI.0003-15.2015.

67. Wei, X., Ma, T., Cheng, Y., Huang, C.C.Y., Wang, X., Lu, J., and Wang, J. (2018). Dopamine D1 or D2 receptor-expressing neurons in the central nervous system. Addict Biol 23, 569–584. 10.1111/adb.12512.

68. West, E.A., Niedringhaus, M., Ortega, H.K., Haake, R.M., Frohlich, F., and Carelli, R.M. (2021). Noninvasive Brain Stimulation Rescues Cocaine-Induced Prefrontal Hypoactivity and Restores Flexible Behavior. Biol Psychiatry 89, 1001–1011. 10.1016/j.biopsych.2020.12.027.

69. Wickersham, I.R., Lyon, D.C., Barnard, R.J., Mori, T., Finke, S., Conzelmann, K.K., Young, J.A., and Callaway, E.M. (2007). Monosynaptic restriction of transsynaptic tracing from single, genetically targeted neurons. Neuron 53, 639–647. 10.1016/j.neuron.2007.01.033.

70. Witten, I.B., Lin, S.C., Brodsky, M., Prakash, R., Diester, I., Anikeeva, P., Gradinaru, V., Ramakrishnan, C., and Deisseroth, K. (2010). Cholinergic interneurons control local circuit activity and cocaine conditioning. Science 330, 1677–1681. 330/6011/1677 [pii]10.1126/science.1193771.

71. Ztaou, S., Oh, S.J., Tepler, S., Fleury, S., Matamales, M., Bertran-Gonzalez, J., Chuhma, N., and Rayport, S. (2021). Single dose of amphetamine induces delayed subregional attenuation of cholinergic interneuron activity in the striatum. eNeuro 8, 1–13. 10.1523/eneuro.0196-21.2021.

72. Zucker, R.S., and Regehr, W.G. (2002). Short-term synaptic plasticity. Annu Rev Physiol 64, 355–405.

